# Impact of vitamin D deficiency on defective endometrial decidualization and the repressive role of vitamin D receptor (VDR) in the epigenomic network

**DOI:** 10.1101/2025.11.14.688535

**Authors:** MyeongJin Yi, Skylar G Montague Redecke, Tianyuan Wang, Austin Bell-Hensley, Shuyun Li, Abdull J. Massri, Anne Marie Z. Jukic, Francesco J. DeMayo

**Affiliations:** Reproductive and Developmental Biology Laboratory, National Institute of Environmental Health Sciences, National Institutes of Health, Research Triangle Park, North Carolina, 27709, United States; Integrative Bioinformatics Supportive Group, National Institute of Environmental Health Sciences, National Institutes of Health, Research Triangle Park, North Carolina, 27709, United States; Epidemiology Branch, National Institute of Environmental Health Sciences, National Institutes of Health, Research Triangle Park, North Carolina, 27709, United States

**Author notes:** Corresponding authors: National Institute of Environmental Health Sciences, National Institutes of Health, Research Triangle Park, North Carolina, 27709, United States. E-mail addresses (F.J. DeMayo), (A.M.Z. Jukic).

**Keywords:** Vitamin D, VDR, decidualization, pregnancy, epigenomics, transcriptomics

## Abstract

Identifying the factors that regulate female reproduction is crucial to understanding how the environment affects female reproductive health. The vitamin D receptor (VDR) and its ligands (primarily 1,25(OH)_2_D_3_) have a recognized role in calcium homeostasis; however, their broader impact on female reproduction remains underexplored. We demonstrate that the VDR and its ligands are involved in the hormonal induction of uterine decidualization. Mice fed a vitamin D-deficient diet displayed an impaired hormonally induced decidual response. In a human telomerase reverse transcriptase-immortalized human endometrial stromal cell line (T-HESC), VDR decreased during in vitro decidualization. Small interfering RNA (siRNA) knockdown of VDR in T-HESC enhanced in vitro decidualization, while overexpression of VDR inhibited it. Chromatin accessibility and histone modification analyses revealed that VDR functions as a chromatin regulator, restricting accessibility and repressing transcription in specific genomic regions. Transcriptomic analyses confirmed that VDR broadly modulates gene expression, with most ligand-mediated effects occurring through the VDR. These findings identify VDR as a key regulator of transcriptional and chromatin landscapes in human endometrial stromal cells, offering novel insights into vitamin D signaling in reproduction. This study highlights the potential of targeting vitamin D pathways to treat uterine disorders associated with impaired decidualization and reduced fertility.

**Graphical Abstract:** 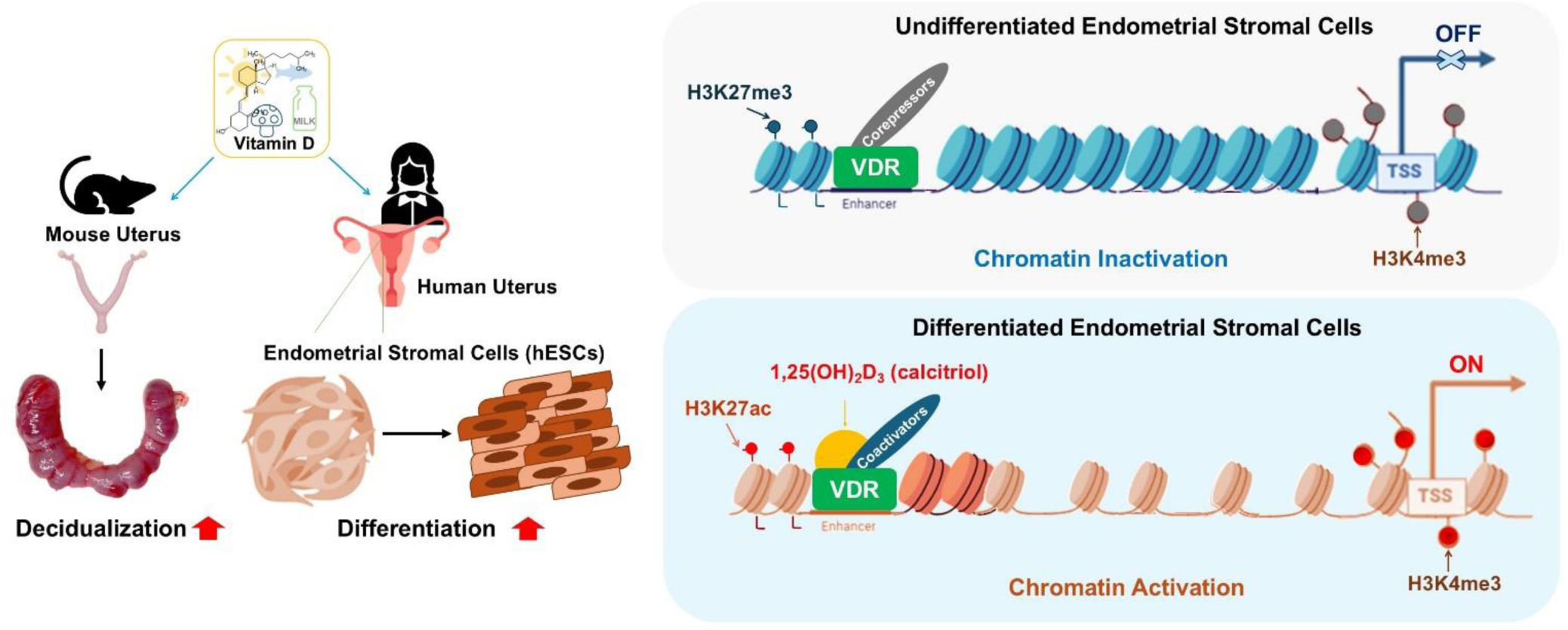

## 1. Introduction

Vitamin D is traditionally associated with calcium metabolism [1] and bone health [2], however, its role in female reproductive health is a promising area of research [3]. Vitamin D, or 1α,25-dihydroxyvitamin D_3_ (1,25(OH)_2_D_3_, calcitriol) in its active form, functions as a potent ligand for vitamin D receptor (*VDR*), modulating the expression of numerous genes involved in cellular differentiation [4, 5], immune regulation [6], and inflammation [7–9]. VDR is expressed in reproductive tissues, including the ovaries [10], endometrium, and placenta [11], highlighting its potential role in reproduction. The association between vitamin D deficiency and adverse reproductive outcomes has garnered significant attention. Epidemiological studies have linked vitamin D deficiency to reduced fertility [12, 13], implantation failure [14–16], and pregnancy complications such as preeclampsia [11, 17, 18] or gestational diabetes [18–21]. Additionally, vitamin D appears to influence ovarian function [22], follicular development, and estradiol across the menstrual cycle [23], further emphasizing its multifaceted role in reproduction. For instance, in women undergoing in vitro fertilization (IVF), higher serum vitamin D levels correlate with improved live birth rates, although some variability exists across studies [24]. Similarly, in women with polycystic ovary syndrome (PCOS), vitamin D supplementation has shown potential benefits [25], including improved endometrial thickness [26] and enhanced ovulatory function [27].

Defective uterine receptivity represents a significant barrier to successful pregnancy, contributing to infertility and recurrent pregnancy loss [28–30]. The intricate molecular events within the female reproductive system are critical for establishing and sustaining pregnancy [31–33]. Among these events, hormonally regulated endometrial decidualization is a critical process by which fibroblast-like endometrial stromal cells transform into specialized secretory decidual stromal cells, creating a conducive uterine environment for embryo implantation [34] and placental development [35, 36]. This process involves hormonal signals, immune modulation, and cellular reprogramming, all orchestrated to prepare the endometrium for pregnancy. Impaired decidualization has been linked to infertility and pregnancy complications, including recurrent miscarriages and uteroplacental disorders [37–39]. While substantial progress has been made in understanding the cellular mechanisms of decidualization and stromal cell differentiation, further research is needed to unravel the regulatory networks and identify novel therapeutic targets for decidualization defects.

VDR and its ligands have been implicated in regulating the processes of decidualization and endometrial stromal cell differentiation [14]. Mice with ablation of the VDR demonstrate uterine hyperplasia and defects in ovarian function [40]. The presence of VDR in the endometrium suggests a role for vitamin D signaling and the hormonal and molecular pathways critical for endometrial receptivity. Recent studies suggest that 1,25(OH)_2_D_3_ influences the expression of key decidualization markers, such as prolactin (PRL) and insulin-like growth factor-binding protein 1 (IGFBP1) [14, 41]. Vitamin D signaling may enhance decidualization, creating a more favorable environment for implantation and early pregnancy.

Our objective was to investigate the role of vitamin D and VDR in endometrial stromal cell differentiation and to elucidate the molecular mechanisms. We leveraged both an in vivo mouse model of dietary vitamin D_3_ deficiency to examine fertility performances and in vitro analyses of human endometrial stromal cells. The findings of this research not only enhance our understanding of the role of the vitamin D pathway in female reproductive physiology but also provide insights into potential therapeutic interventions for addressing infertility and pregnancy-related disorders.

## 2. Materials and methods

### 2.1. Materials

Key sources used in this study are detailed in the supplementary material table 1. The table includes information on reagent types, resources, compound names, manufacturers, references, and identifiers such as catalog numbers and original distributors.

### 2.2. Mice

C57BL/6J mice were obtained from The Jackson Laboratory. Animals were fed either a vitamin D_3_-replete control diet (TD.10415, Teklad) or a vitamin D_3_-deficient diet (TD.09116, modified from TD.89123, Teklad). Detailed nutrient composition is provided in supplementary material table 1. Mice were housed under standard conditions at the Comparative Medicine Branch (CMB), and all animal procedures were conducted in accordance with protocols approved by the NIEHS/NIH Animal Care and Use Committee (ACUC). All methods complied with Public Health Service Policy and included humane examinations to ensure animal welfare.

### 2.3. Justification of animal numbers

For the animal experiments, the sample size for each group was determined based on the power analysis. Assuming a 1:1 control-to-case ratio, if the hypothetical portion with exposure (vitamin D deficiency) to the case is regarded as 99, and the control is regarded as 1, five mice per group will be sufficient to observe with a power of 80% chance of detecting, and two-sided confidence level (1-α) = 95, based on Kelsey and Fleiss with a continuity correction [42]. For a scenario where the hypothetical exposure proportion (vitamin D deficiency) to the case is regarded as 99.99, and the control is 0.01, a minimum of four mice per group is sufficient for a single comparison.

### 2.4. Artificial decidualization

Female mice, 6 weeks old at the start of dietary intervention, were fed vitamin D control or deficient diets for ∼5 weeks to prime total serum 25-hydroxyvitamin D (25(OH)D) levels. Mice were ovariectomized at 11 weeks of age and allowed 2 weeks for hormone stabilization. To induce artificial decidualization, mice received daily subcutaneous injections of 100 ng 17β-estradiol for 3 consecutive days, followed by a 2-day rest [43, 44]. This was followed by daily injections of 1 mg progesterone and 6.7 ng 17β-estradiol for 3 days [43, 44]. On the third day of progesterone/estradiol treatment, 50 μL sesame oil was injected into one uterine horn 6 h after hormone administration. Mice continued daily progesterone/estradiol injections for 5 additional days, and uteri were collected on day 6 [43, 44]. Both oil-injected and contralateral horns were weighed (N = 9). Across priming, stabilization, and hormone treatment, mice received the assigned diet for approximately 9 weeks. For ovariectomy and oil injection procedures, mice received a pre-operative analgesic 10-30 min prior to anesthesia. Anesthesia was induced via inhalation of isoflurane delivered through a calibrated vaporizer to minimize pain and stress, and a toe-pinch reflex test was performed prior to all surgical procedures to confirm a fully unconscious state.

### 2.5. Cell culture of human telomerase reverse transcriptase (hTERT)-immortalized human endometrial stromal cell line (T-HESC)

T-HESC was purchased from the American Type Culture Collection (ATCC). The cells were cultured and maintained in Dulbecco’s Modified Eagle Medium/Nutrient Mixture F-12 (DMEM/F12), which was supplemented with 10% (v/v) fetal bovine serum (FBS), 1 mM sodium pyruvate, 100 units/mL penicillin, and 100 µg/mL streptomycin at 37°C, containing 5% CO_2_ in a humidified atmosphere [45–47]. All experiments were performed using T-HESC between passages 25 and 30.

### 2.6. siRNA transfection

T-HESCs were transfected with nontargeting siRNA (siNT; ON-TARGETplus Non-targeting Control Pool, Dharmacon) or siRNA targeting VDR (siVDR; ON-TARGETplus SMARTpool, Dharmacon) using Lipofectamine RNAiMAX (Invitrogen) according to the manufacturer’s protocol. Briefly, 10⁵ cells per well were seeded in a 6-well plate and cultured for 24 hr in standard endometrial stromal cell medium. For each well, 5 μL of Lipofectamine RNAiMAX was diluted in 145 μL of Opti-MEM in a sterile microtube, and 3 μL of 10 μM siRNA was diluted in 97 μL of Opti-MEM in a separate microtube. The diluted solutions were incubated individually for 5 min at room temperature, then combined in the same tube and incubated for an additional 10 min to allow complex formation. The resulting siRNA-lipid complexes (250 μL) were added dropwise to each well, followed by 1 mL of Opti-MEM supplemented with 2% (v/v) charcoal-stripped FBS (antibiotic-free). Plates were gently swirled to ensure even distribution, and cells were incubated for 48 hr at 37 °C in a humidified 5% CO₂ atmosphere before downstream analyses.

### 2.7. Lentiviral transduction

A hemagglutinin (HA)-tagged human VDR lentiviral expression plasmid and a green fluorescent protein (GFP)-inserted lentiviral control vector were designed and purchased from Applied Biological Materials (ABM). The lentivirus plasmid was packaged by the Viral Core at NIEHS. T-HESCs were transduced with either the HA-tagged VDR plasmid or the control plasmid in the presence of polybrene at a multiplicity of infection (MOI) of 10. Cells were incubated for 24 hr in the presence of lentivirus before changing the media. The successfully transfected cells were selected by puromycin antibiotic-containing culture medium for an additional 24 hr.

### 2.8. Exogenous hormonal exposure for in vitro decidualization

For in vitro decidualization, T-HESCs were seeded into a 6-well plate at a density of 10^5^ cells/well in normal culture medium. After incubating the cells for 24 hr, the medium was discarded, washed with 1 mL of Opti-MEM, and the medium was replaced with either decidual or vehicle medium. The decidual medium contained Opti-MEM supplemented with 2% (v/v) charcoal-stripped FBS, 1 mM sodium pyruvate, 100 units/mL penicillin, 100 µg/mL streptomycin, 10 nM 17β-estradiol, 1 µM medroxyprogesterone acetate (MPA), and 100 µM dibutyryl cyclic adenosine monophosphate (db-cAMP) sodium salt, collectively referred to as EPC. This treatment induces morphological changes and upregulates the expression of decidual markers, prolactin (*PRL),* and insulin-like growth factor-binding protein 1 (*IGFBP1*) production [48–53]. The vehicle medium contained Opti-MEM supplemented with 2% charcoal-stripped FBS (v/v), 1 mM sodium pyruvate, 100 units/mL penicillin, 100 µg/mL streptomycin, and 200 proof ethyl alcohol at a volume that corresponded with the volume of exogenous hormonal stock used. The total hormonal exposure time was 72 hr, and the corresponding medium was replaced after 48 hr.

### 2.9. Quantitative real-time PCR (qRT-PCR)

Total RNAs from T-HESCs were isolated using TRIzol reagent according to the manufacturer’s instructions. All isolated RNAs were quantified and certified using a Qubit RNA High Sensitivity kit (Invitrogen). Each cDNA was synthesized with a High-Capacity cDNA Reverse Transcription kit (Applied Biosystems), adding 2 µg of total RNA to the reaction mixture, and the reaction mixtures were incubated at room temperature for 1 hr followed by additional incubation at 37°C for 2 hr. Each synthesized cDNA was quantified and certified using a Qubit dsDNA High Sensitivity kit (Invitrogen). Taqman probes for human genes were used as Hs99999901_s1, Hs00236877_m1, Hs00168730_m1, and Hs00172113_m1 for 18*s, PRL*, *IGFBP1*, and *VDR*, respectively. The probes for mouse genes were used as Mm01340178_m1, Mm01194003_m1, Mm00494566_m1, and Mm00435625_m1 for *Bmp2*, *Wnt4*, *Prl8a2*, and *Pgr*, respectively. qPCR was performed with the CFX96 Real-Time PCR Detection System (Bio-Rad). Each value was derived from the comparative CT method, which compared the *C_t_* value of one target gene to a reference gene using the 2^-ΔΔ*Ct*^ formula according to the manufacturer’s guidelines. Δ*C_t_* indicates the differences in threshold cycles for target and reference (*C_t_*,_target_ – *C_t_*,_reference_), and ΔΔ*C_t_* represents the relative change in these differences between the target and reference (Δ*C_t_*,_target_ – Δ*C_t_*,_reference_). Therefore, the expression of the target, normalized to a housekeeping gene, was given by 2^-ΔΔ*Ct*^.

### 2.10. Western blot analysis

Total protein lysates from T-HESCs were isolated using RIPA lysis buffer. Cells were washed three times with cold PBS, after which 500 μL of RIPA buffer was added directly to each well and lysates were collected with a cell scraper. The suspension was transferred to pre-chilled microtubes. As a positive control, kidney lysates were prepared from two female C57BL/6J mice. Approximately 50 mg of kidney tissue was washed with PBS and homogenized using a Dounce homogenizer in 1 mL of RIPA buffer. T-HESC and kidney lysates were vortexed for 15 s, incubated on ice for 5 min, and centrifuged at 12,000 × g for 15 min. Supernatants were transferred to pre-chilled tubes, and protein concentrations were determined using the Bio-Rad Protein Assay Dye Reagent. Due to the high VDR abundance in the kidney, 15 μg of kidney lysate and 30 μg of T-HESC lysate were loaded per lane. Samples were separated on a 10% SDS-PAGE gel and transferred onto a PVDF membrane. Membranes were blocked in 5% (w/v) BSA in TBS containing 0.1% (v/v) Tween-20 (TBS-T). HRP signals were detected using an ECL kit and imaged using a Bio-Rad ChemiDoc system.

### 2.11. Assessment of chromatin accessibility of VDR by ATAC-seq

T-HESCs were transfected with either siVDR or siNT, and nuclei were extracted for library preparation using previously published procedures [54, 55]. The cells were from three technical replicates, and in total, 6 samples were collected for ATAC-seq analysis according to the published protocol [54, 55]. Briefly, cells were extracted using Greenleaf buffer, containing 10 mM Tris-HCl, 10 mM sodium chloride, 3 mM magnesium chloride, and 0.1% IGEPAL CA-630, and homogenized in a chilled douncer. Nuclei were counted using a hemacytometer, and 50,000 nuclei were used in the transposition reaction as described previously [54]. Sequencing was performed on a NovaSeq (Illumina). Raw reads (50 bp, paired-end) were processed by trimming adaptors and filtering with average quality scores greater than 20 by Trim Galore version 6.7. The reads passing the initial processing were aligned to the human reference genome (hg38) using bowtie with unique mapping and up to 2 mismatches for each read (-m 1 -v 2). After removing the reads mapped to mitochondrial DNA and the duplicated reads, the uniquely mapped duplicated reads in each sample were normalized by down-sampling to 100 million. Only the first 9 bp of each read were used for downstream analyses. The open chromatin regions were first identified by MACS2 with a cutoff of Q-value 0.0001, followed by merging genomic intervals within 100 bp of each other [56, 57].

### 2.12. Analyses of histone modification markers and distribution of chromatin-associated proteins by CUT&RUN

CUT&RUN samples were prepared with the CUTANA^TM^ ChIC/CUT&RUN kit (EpiCypher), following the manufacturer’s guidelines with modifications. The libraries were sequenced by the NIEHS Sequencing Core. The reads were trimmed by adapter sequences using Trim Galore, aligned to hg38 and E. coli K-12 MG1655 genomes using Bowtie2 (v2.5.2), deduplicated using Picard, and filtered using samtools. MACS2 was used to call peaks for each sample; peaks were combined with all triplicates of each condition, and the peaks were used for further analyses [56–59]. All peaks were annotated for overlaps with transcriptome and chromatin assessment analyses.

### 2.13. Transcriptomic analysis by bulk RNA-seq

The library was prepared using the TruSeq RNA Library Prep kit (Illumina) and subsequently sequenced using NextSeq 500. The sequencing reads with a quality score < 20 were filtered using a custom Perl script [60]. The adaptor sequence was removed using Cutadapt (v1.12). The reads were aligned to the hg38 genome using the STAR aligner (v2.5.2b) and counted using the featureCounts (v1.5.0-p1) function in the Subread program. The differentially expressed genes (DEG) between siNT and siVDR, or vehicle- and 1,25(OH)_2_D_3_-treated, were identified using Partek Genomic Suite [61]. The threshold was set as “maximal FPKM ≥ 1, false discovery rate (FDR) adjusted P < 0.05, fold change ≥ 1.4 (up-regulated) or ≤−1.4 (down-regulated).”

### 2.14. Annotation of the nearest gene and motif analysis

The gene associated with each peak was predicted by searching the transcription start site (TSS) of nearby genes within a 100 kb range using the ‘annotatePeaks.pl’ function of the Hypergeometric Optimization of Motif Enrichment (HOMER) motif discovery tool [62]. HOMER’s findMotifsGenome.pl function was used for motif enrichment analysis of given peak ranges.

### 2.15. Statistical analysis

All data were first tested for normality. If the normality assumption was met, comparisons were done using one-way ANOVA with Tukey’s multiple comparisons for multiple groups and a Student’s *t*-test for two groups. For non-normal distributed data, the Kruskal-Wallis test with Dunnett’s multiple comparisons was used for multiple groups, and the Mann-Whitney test was used for two groups.

## 3. Results

### 3.1. Vitamin D deficiency impairs uterine decidualization in mice

Previous studies in global VDR knockout mice demonstrated abnormalities in bone mineral homeostasis and slight hypocalcemia, accompanied by compensatory increases in circulating 1,25(OH)_2_D_3_ [40, 63]. These mice also showed uterine hyperplasia and impaired folliculogenesis. To further investigate the role of vitamin D in uterine decidualization, we induced vitamin D deficiency in 6-week-old female C57Bl/6J mice by feeding them either a vitamin D_3_-sufficient diet (1000 IU/kg) or a vitamin D_3_-deficient diet (0 IU/kg) for nine weeks, including hormonal injection periods (Fig. 1A). The total serum 25(OH)D levels were significantly altered after at least six weeks of diet feeding and almost abolished at eight weeks of diet (Supplementary material figure 1A). We then investigated reproductive parameters, including the estrous cycle, superovulation (Supplementary material figure 1B), and long-term fertility over a 6-month breeding trial (Supplementary material figure 1C). For the estrous cycle analysis, mice were primed with their respective diets for 8 weeks before tracking cycle progression over 15 days. There were no obvious disruptions in the estrous cycle of vitamin D-deficient mice compared to controls (Supplementary material figure 2A). Similarly, superovulation was assessed by counting the number of oocytes collected after hormonal stimulation. No significant differences in oocyte numbers were observed between the diet groups, although the sample size was small (Supplementary material figure 2B).

**Fig. 1.**
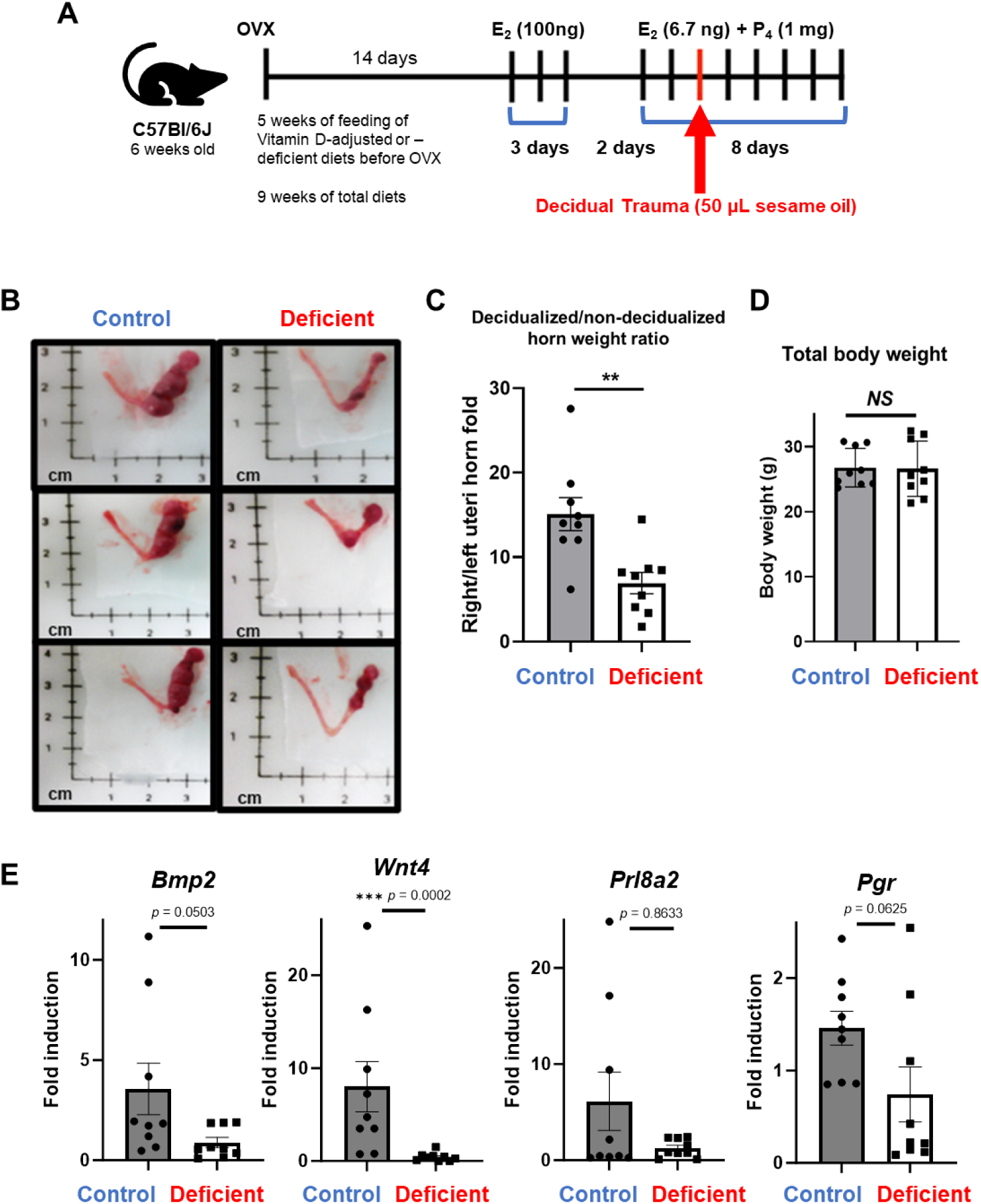
Impaired decidualization in vitamin D-deficient mice. (A) Experimental schematic for inducing vitamin D deficiency in female C57Bl/6J mice through dietary intervention and subsequent artificial decidualization. (B) Representative images of uterine horns after decidualization. (C) Quantification of the weight ratio of oil-injected uterine horns to uninjected horns. (D) Body weights of vitamin D-sufficient and vitamin D-deficient mice. (E) qRT-PCR analysis of decidual marker gene expressions in uterine tissue, such as *Bmp2*, *Wnt4*, *Prl8a2*, and *Pgr*. Data are presented as mean ± SEM. Statistical significance: *p < 0.05, **p < 0.01, ***p < 0.001.

To evaluate long-term reproductive outcomes, we conducted a 6-month breeding trial. Since the timeline of the trial was extended, dietary priming was not performed prior to mating (Supplementary material figure 1C). Fertility indices, including the total number of litters and pups, were comparable between the vitamin D-sufficient and vitamin D-deficient groups (Supplementary material figure 2C). However, a downward trend in the cumulative number of pups born to vitamin D-deficient dams was observed (Supplementary material figure 2D). Moreover, three vitamin D-deficient dams were found deceased during their first littering event, likely due to dystocia, a condition characterized by impaired muscle contraction during delivery (Supplementary material figure 2E).

Given the trend towards decreased fertility in the vitamin D-deficient diet group, we next investigated the response of the uterus to a hormonally induced differentiation, the decidual assay. Female mice on a control of a vitamin D-deficient diet were ovariectomized and then given a regimen of estradiol and progesterone, followed by injection of oil into one uterine horn to induce the decidual response. The contralateral uninjected horn served as a control. [64, 65]. In control mice, the weight ratio of the oil-injected uterine horn to the uninjected horn increased significantly (15.09-fold ± 1.941 SEM). However, this response was impaired in vitamin D-deficient mice, as shown by a reduced weight ratio compared to the control (6.917-fold ± 1.259 SEM) (Fig. 1B and C). Body weight measurements revealed no differences between the two groups, ruling out systemic effects of the diet on overall health (Fig. 1D). To further explore the molecular impact of vitamin D deficiency on uterine function, we examined the expression of decidual marker genes, including *Bmp2*, *Wnt4*, *Prl8a2*, and *Pgr,* using qRT-PCR. All tested genes showed reduced expression in vitamin D-deficient mice, with *Wnt4* displaying a statistically significant decrease (Fig. 1E). These results indicate that vitamin D deficiency compromises both the morphological and molecular aspects of uterine decidualization.

Taken together, these findings demonstrate that while vitamin D deficiency does not significantly alter estrous cycling or superovulation, it has a profound impact on uterine decidualization. This disruption may contribute to increased reproductive risks, including complications during labor, under conditions of vitamin D deficiency.

### 3.2. Vitamin D receptor expression is suppressed during in vitro decidualization of telomerase-transformed human endometrial stromal cells, T-HESC

Building on the mouse study indicating vitamin D’s supportive role in decidualization, we investigated its role in human in vitro decidualization using the immortalized human endometrial stroma cell line, T-HESC. Upon treatment of T-HESCs in decidualization media containing EPC for 72 hr, *VDR* mRNA levels were significantly reduced (Fig. 2A). Western blot analysis confirmed decreased VDR protein levels in EPC-treated cells, supporting VDR suppression in differentiated stromal cells (Fig. 2B). To further elucidate VDR’s role, we transfected T-HESCs with siRNA targeting VDR (siVDR) or a non-targeting control (siNT). siVDR significantly reduced VDR mRNA levels (Fig. 2C). EPC treatment further decreased *VDR* expression in both siNT and siVDR cells. Notably, VDR knockdown led to a marked increase in *PRL* and *IGFBP1* production during differentiation compared to controls (Fig. 2D and E). These results indicate that a decrease in VDR is a requisite for full endometrial stromal cell decidualization.

**Fig. 2.**
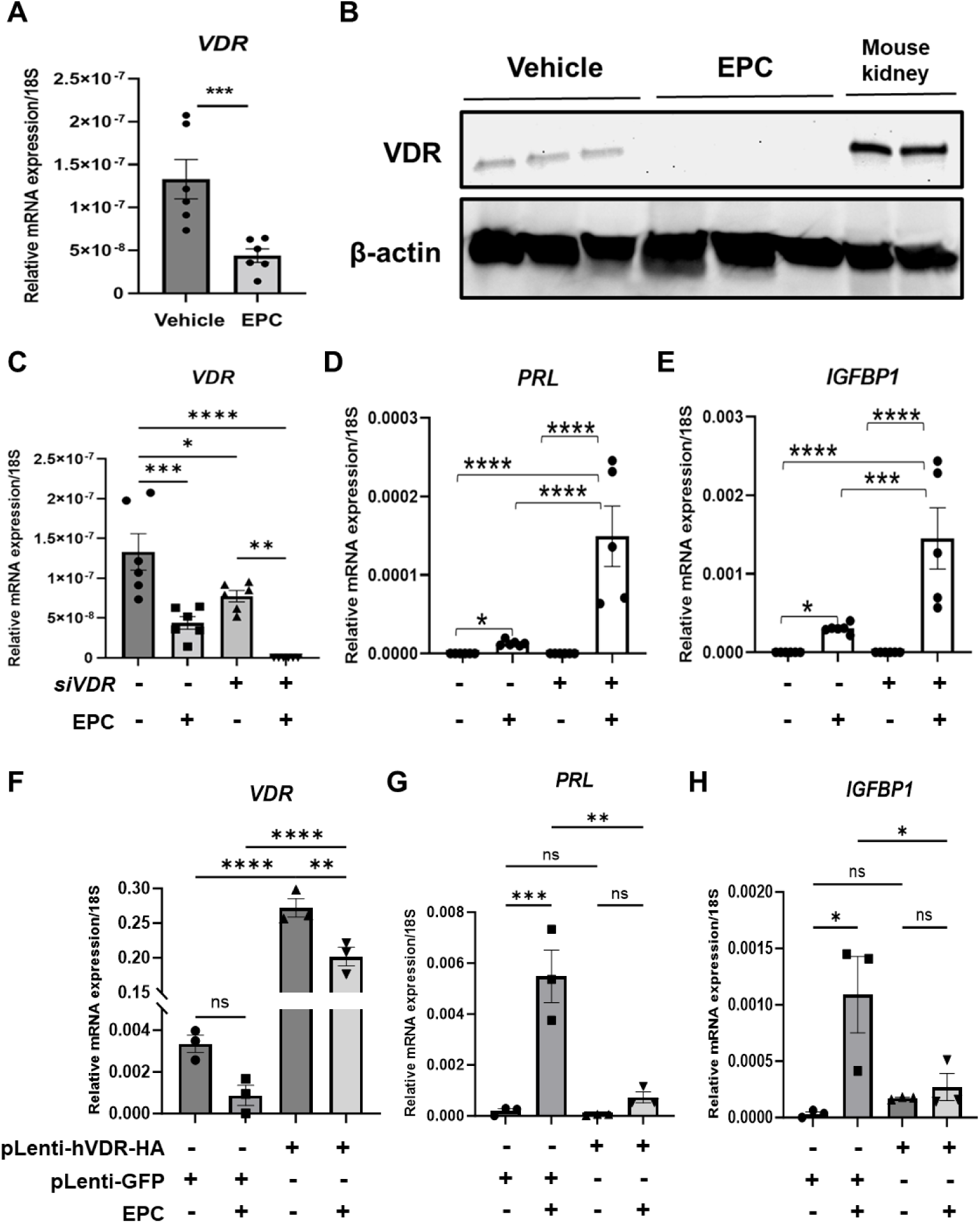
VDR suppression during hormone-induced decidualization of human endometrial stromal cells (T-HESCs). (A) VDR mRNA expression levels in T-HESCs treated with EPC compared to untreated controls. (B) Western blot analysis of VDR protein levels in T-HESCs over 72 hr of EPC treatment, with kidney lysates from two individual female C57BL6/J mice as a positive control. (C) Validation of siVDR knockdown efficiency by qRT-PCR. (D) PRL and (E) IGFBP1 mRNA expression in siNT- and siVDR-transfected cells during EPC treatment. (F) VDR overexpression in lentiviral transduced T-HESCs was confirmed by qRT-PCR, and pLenti-GFP vector transduced T-HESCs were used as a control. (G) PRL and (H) IGFBP1 mRNA expression in VDR-overexpressing cells during EPC treatment. Data are presented as mean ± SEM. Statistical significance: *p < 0.05, **p < 0.01, ***p < 0.001, ****p < 0.0001.

In order to determine whether a decrease in VDR is required for decidualization, VDR expression was maintained throughout decidualization by utilizing lentiviral transduction with HA-tagged VDR plasmids. A virus expressing GFP-only vectors served as a control. Lentiviral transduction significantly increased *VDR* mRNA levels compared to GFP controls (Fig. 2F). In alignment with the knockdown results, VDR overexpression significantly reduced *PRL* and *IGFBP1* mRNA levels during differentiation (Fig. 2G and H). These findings demonstrate that VDR functions as a suppressor of human endometrial stromal cell differentiation, and its downregulation is necessary for full decidualization.

### 3.3. VDR knockdown enhances chromatin accessibility and alters histone modifications in open chromatin regions in T-HESCs

To further investigate the suppressive role of VDR during endometrial stromal cell differentiation, we assayed the impact of VDR knockdown on open chromatin accessibility and histone modification. ATAC-seq analysis was conducted to evaluate the impact of VDR knockdown on chromatin accessibility in T-HESCs. In cells transfected with control siRNA (siNT), a total of 45,230 open chromatin peaks were identified (Supplementary material table 2). In contrast, cells with VDR knockdown (siVDR) displayed a significant increase, with 73,238 peaks observed (Supplementary material table 2). Altogether, 78,972 unique peaks were identified across both conditions, of which 39,496 peaks were shared between siNT and siVDR. These shared peaks accounted for 87.3% of the siNT peaks and 53.9% of the siVDR peaks (Fig. 3A). This substantial overlap indicates that while a core set of chromatin regions remains accessible regardless of VDR presence, the loss of VDR induces the accessibility of additional regions, leading to a broader chromatin accessibility (Fig. 3B).

**Fig. 3.**
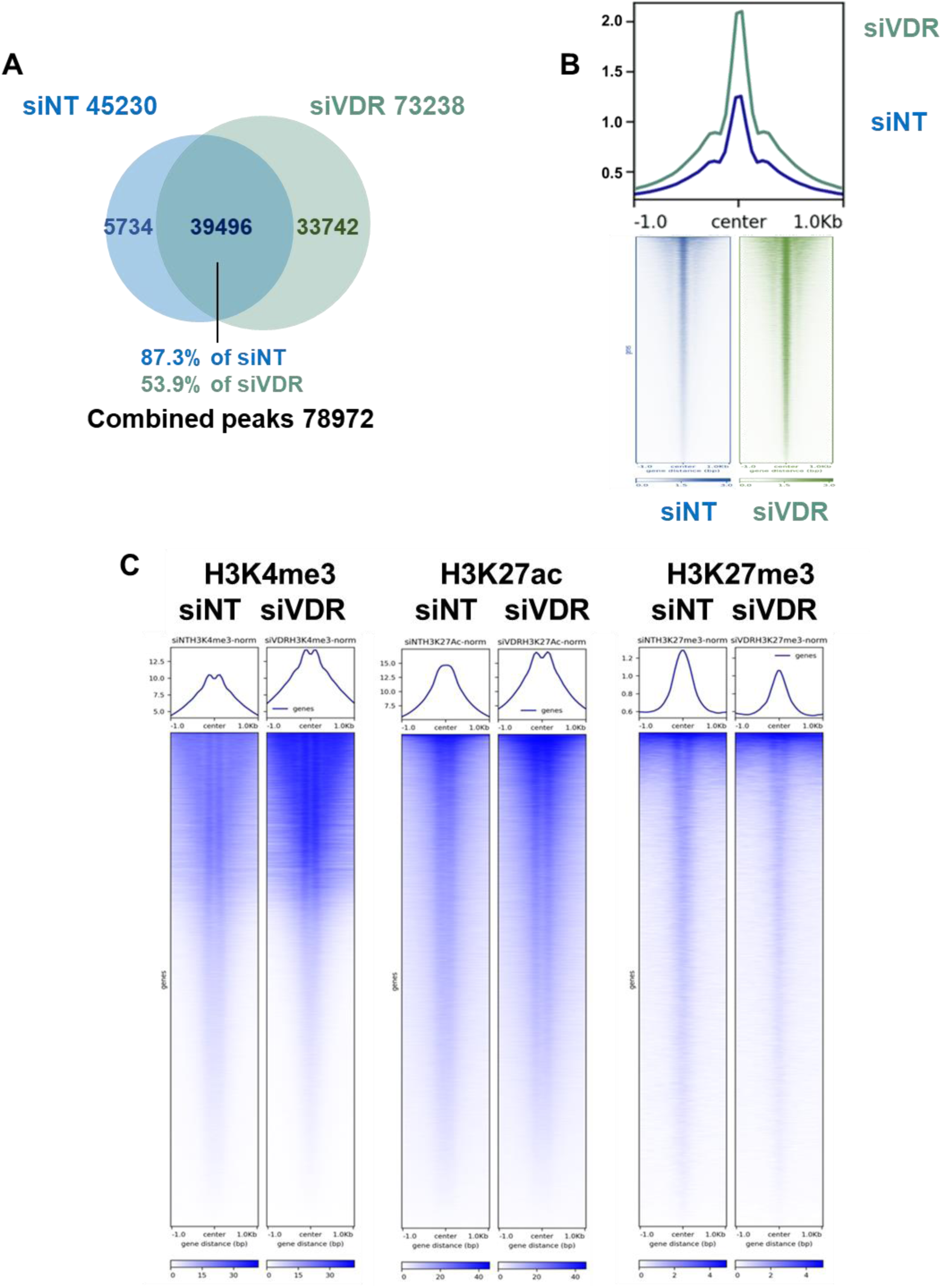
Chromatin accessibility and histone modifications in VDR knockdown T-HESCs. (A) Overlap of open chromatin peaks identified by ATAC-seq in siNT and siVDR cells. (B) Chromatin accessibility across universal open chromatin regions is plotted as metaplots and heatmaps. The ATAC-seq reads from all technical replicates are combined. (C) Comparison of histone modifications (H3K4me3, H3K27ac, H3K27me3) using CUT&RUN between siNT and siVDR within open chromatin regions. Data indicate increased transcriptionally active marks (H3K4me3, H3K27ac) and decreased repressive marks (H3K27me3) in siVDR-transfected cells.

To further investigate the relationship between VDR and chromatin regulation, we profiled histone modifications genome-wide using CUT&RUN. We then examined how these marks were distributed both globally and within the 78,972 open chromatin peaks identified by ATAC-seq. In siVDR cells, the levels of H3K4me3 and H3K27ac, active histone marks associated with promoters and enhancers, were markedly increased compared to controls (Fig. 3C). This indicates enhanced transcriptional activity in regions where chromatin became more accessible following VDR knockdown. Conversely, the levels of the repressive histone mark H3K27me3 were reduced in siVDR cells, suggesting a decrease in transcriptional repression (Fig. 3C).

Together, these findings reveal that VDR plays a critical role in chromatin regulation by modulating both accessibility and histone modification patterns. The depletion of VDR leads to increased chromatin openness, shifting the balance of histone modifications toward a transcriptionally active state. The role of VDR as a regulator of chromatin accessibility and epigenetic marks underscores its importance in maintaining T-HESCs in human endometrial stromal cell differentiation.

### 3.4. Transcriptomic changes regulated by VDR and 1,25(OH)₂D₃ in T-HESCs

To further investigate the role of VDR in decidualization, we investigated the impact of the VDR ligand, 1,25(OH)₂D₃ in T-HESCs. T-HESCs transfected with siNT or siVDR were treated with EPC and either vehicle or 1,25(OH)₂D₃. PRL, a decidualization marker, was upregulated in response to VDR knockdown under vehicle and 1,25(OH)₂D₃, with 1,25(OH)₂D₃ further enhancing expression (Fig. 4A). These findings align with our in vivo mouse diet studies, supporting a dual role for vitamin D and VDR in regulating stromal cell function during decidualization.

**Fig. 4.**
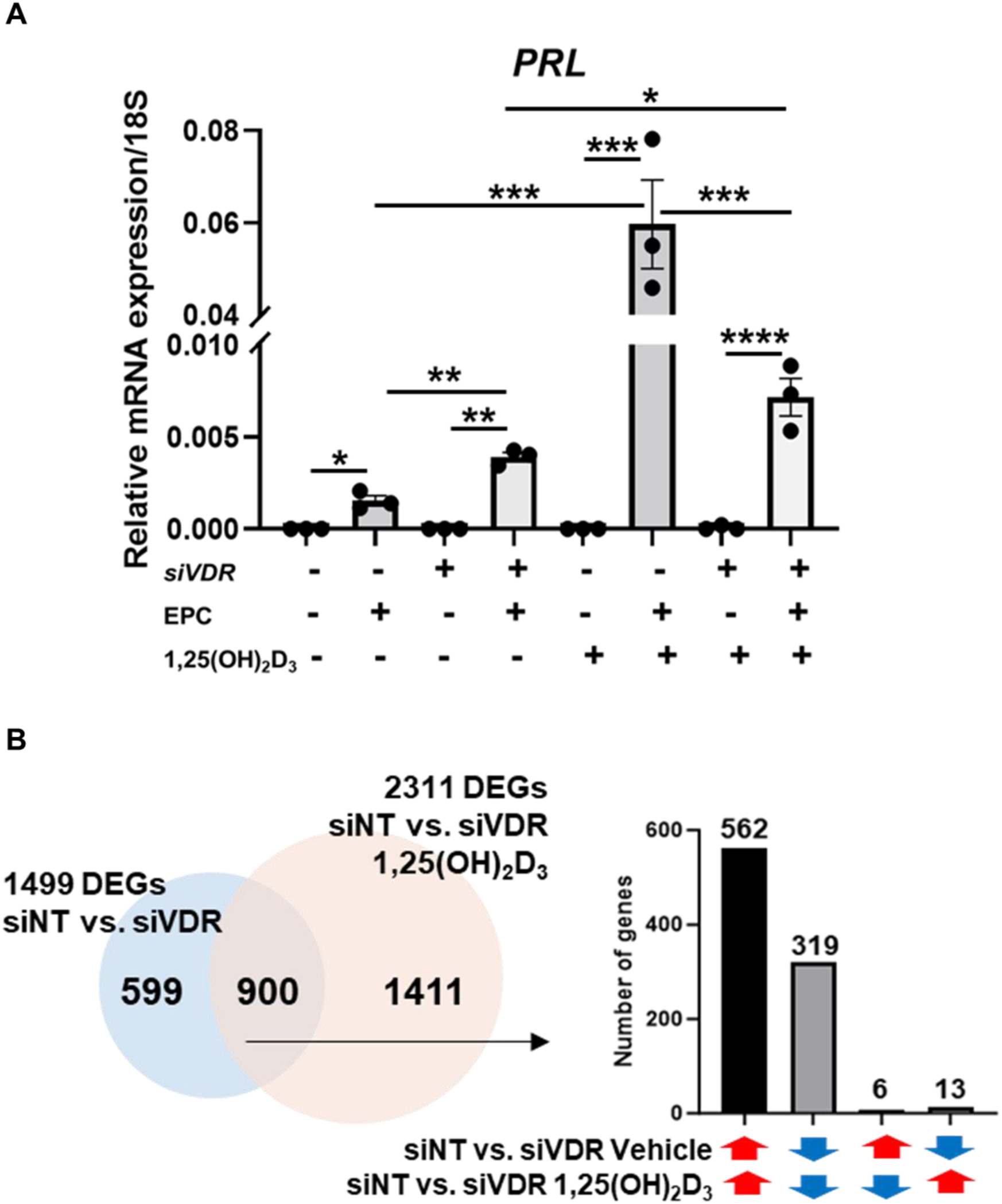
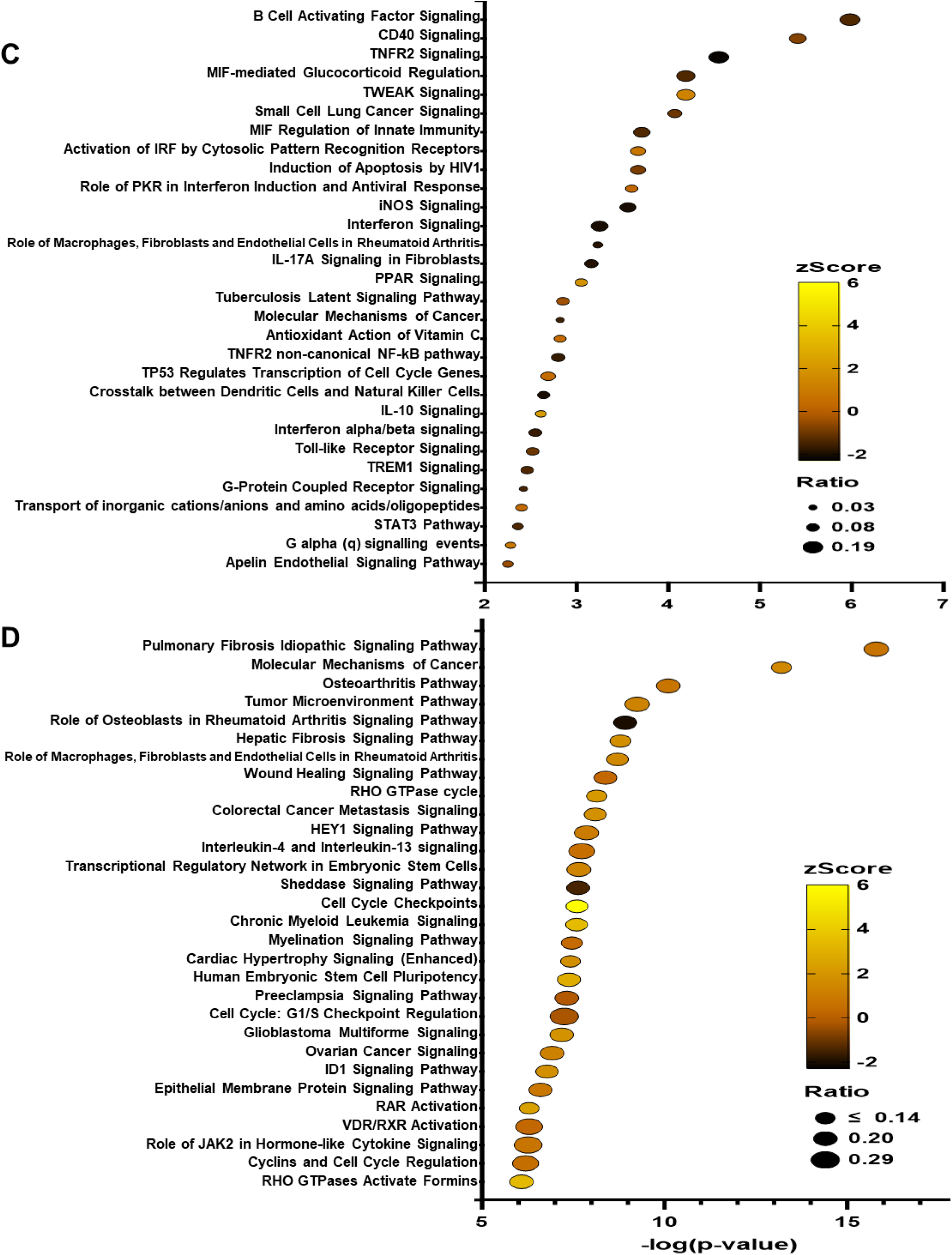
Transcriptomic alterations in T-HESCs by VDR knockdown and 1,25(OH)₂D₃. (A) Prolactin (PRL) expression as a marker of decidualization in T-HESCs transfected with siNT or siVDR and treated with EPC and either vehicle or 1,25(OH)₂D₃. PRL expression increases with 1,25(OH)₂D₃ treatment, but VDR knockdown attenuates decidualization despite ligand presence. (B) Venn diagram showing the overlap of DEGs between siNT vehicle vs. siVDR vehicle and siNT 1,25(OH)₂D₃ vs. siVDR 1,25(OH)₂D₃ comparisons. The 900 shared DEGs indicate a core transcriptional network regulated by VDR and its ligand, with consistent expression directionality. (C) Canonical pathways in transcriptomic changes by the 599 ligand-independent DEGs and the (D) ligand-receptor (siNT 1,25(OH)₂D₃ vs. siVDR 1,25(OH)₂D₃) DEGs.

To delineate the transcriptomic effects of VDR and its ligand, 1,25(OH)₂D₃, on human endometrial stromal cells, RNA-seq was conducted in T-HESCs. Cells were transfected with control siRNA (siNT) or VDR-targeting siRNA (siVDR) for 48 hr, followed by treatment with either vehicle or 2 nM 1,25(OH)₂D₃ for 24 hr. Differentially expressed genes (DEGs) analysis revealed distinct transcriptomic shifts across conditions as indicated by principal component analysis (PCA) (Supplementary material figure 3A) and heatmap (Supplementary material figure 3B).

The results of RNA-seq are summarized in the supplemental material table 3. *VDR* knockdown alone (vehicle-treated siNT vs. siVDR) resulted in 1,499 DEGs (818 upregulated, 681 downregulated), underscoring the impact of unliganded VDR. Treatment of T-HESC with 1,25(OH)₂D₃ (siNT 1,25(OH)₂D₃ vs. siNT vehicle) resulted in 626 DEGs (386 upregulated and 240 downregulated). When siVDR cells were treated with 1,25(OH)₂D₃ (siVDR 1,25(OH)₂D₃ vs. siVDR vehicle) displayed only 56 DEGs, suggesting that the ligand’s transcriptomic effects are predominantly mediated through VDR (Supplementary material figure 3C). The largest number of DEGs was observed in the comparison of T-HESCs treated with siNT and 1,25(OH)₂D₃ versus T-HESCs treated with siVDR and 1,25(OH)₂D₃. This resulted in 2,311 DEGs and indicates the full regulation of the T-HESCs by *VDR* and its ligand (Supplementary material figure 3C and Supplementary material table 3).

To further explore this relationship, we analyzed chromatin accessibility and its impact on transcriptional regulation. By annotating genes from open chromatin regions (Fig. 3A), we identified 19,189 genes. Remarkably, 80.59% of receptor DEGs (1,208/1,499) and 77.97% of ligand DEGs (1,802/2,311) were localized within these open chromatin regions (Supplementary material figure 3D), reinforcing the importance of chromatin accessibility in VDR-mediated transcriptional regulation.

To validate RNA-seq findings, we examined representative genes. *CYP24A1*, a well-established VDR target, was significantly induced by 1,25(OH)₂D₃ in siNT cells but attenuated in siVDR cells, emphasizing the ligand-receptor interplay (Supplementary material figure 3E). *VDR* knockdown also increased *IGFBP1* expression, particularly in vehicle-treated cells, consistent with VDR’s suppressive role in decidualization (Supplementary material figure 3E). *EFL1*, a potential candidate for VDR-ligand regulation, was significantly induced by 1,25(OH)₂D₃ but showed diminished expression in siVDR cells (Supplementary material figure 3E). Similarly, *VEGFA*, which is regulated by vitamin D in a context-dependent manner, was upregulated upon 1,25(OH)₂D₃ treatment, corroborating prior findings on its vitamin D response element (VDRE)-mediated transcriptional control (Supplementary material figure 3E).

### 3.5. Receptor- or ligand-dependent, and ligand-independent transcriptomic regulations in T-HESCs

Given the relationship between VDR and its ligand 1,25(OH)₂D₃, we next analyzed the overlap between the largest DEG datasets (Supplementary material figure 3C): siNT vehicle vs. siVDR vehicle (1,499 DEGs) and siNT 1,25(OH)₂D₃ vs. siVDR 1,25(OH)₂D₃ (2,311 DEGs). A total of 900 DEGs were shared between the two datasets, accounting for 60.04% of the siNT vehicle vs. siVDR vehicle DEGs (Fig. 4B). Particularly, 97.89% of these shared DEGs exhibited consistent directional changes, highlighting a coordinated regulatory mechanism (Fig. 4B). This comparison identified 599 genes from the receptor-dependent dataset that do not overlap with the ligand-dependent set, which means that these 599 genes represent the ligand-independent genes regulated by VDR. We next compared the pathways regulated by the ligand-independent and ligand-dependent data sets to determine the mechanisms by which VDR regulates endometrial stroma biology, and a summary of the pathways identified from the ligand-independent and ligand-dependent DEGs is shown in Fig. 4C and D. Notably, both receptor and ligand DEGs highlighted pathways linked to bone-related signaling and calcium regulation, underscoring the established roles of VDR and 1,25(OH)₂D₃ in calcium absorption. A significant number of the 599 ligand-independent DEGs-associated canonical pathways were related to suppressing biological processes such as immune responses (e.g. B cell activating factor signaling, tumor necrosis factor receptor 2 (TNFR2) signaling, macrophage migration inhibitory factor (MIF) regulation of innate immunity, interferon signaling, etc) (Fig. 4C and Supplementary material table 3). In contrast, the 2,311 ligand-receptor DEG-associated canonical pathways were related to activating immune responses, including pathways involving macrophages, fibroblasts, and endothelial cells in rheumatoid arthritis, wound healing signaling, interleukin-4 and -13, and preeclampsia (Fig. 4D and Supplementary material table 3). Predicted upstream regulators of receptor DEGs included TNF, lipopolysaccharide (LPS), β-estradiol, TGFB1, HGF, CG, MPA, IL1B, IFNG, and 8-bromo-cAMP (Supplementary material table 3). To further specify ligand-dependent and -independent transcriptomic regulation, we analyzed the pathways of the 900 overlapping genes between the 1,499 receptor DEGs (siNT vs. siVDR) and the 2,311 ligand-receptor DEGs (siNT 1,25 vs. siVDR 1,25) (Fig. 4B). While ligand-independent DEGs were mostly associated with suppressed canonical pathways (Fig. 4C), the 900 overlapping genes exhibited activation of multiple pathways, including pulmonary fibrosis idiopathic signaling, molecular mechanisms of cancer, and chronic myeloid leukemia signaling (Supplementary material figure 4). Upstream regulators such as β-estradiol, MPA, TGFB1, and LPS were predicted to activate this shared DEG set (Supplementary material table 3). This divergence suggests that although the ligand (1,25(OH)₂D₃) exerts its effects through the receptor (VDR), the ligand may mitigate the receptor’s suppressive role.

Examination of the pathways regulated by the ligand-independent and ligand-dependent pathways shows pathways involved in inflammation, cell growth, and metabolism. To dissect the specific contributions of ligand and receptor, we compared the directionality of canonical pathways and upstream regulators from the ligand-independent and ligand-dependent pathways. The ligand-independent DEGs were linked to 84 pathways, while the ligand-receptor DEGs were associated with 331 canonical pathways (Fig. 5A). Among the 43 overlapping pathways, only 13 displayed consistent directional changes (i.e., activated or inhibited) in both datasets (Fig. 5A). Notably, pathways related to “Inflammation and Immune Response” (e.g., TNFR2 signaling, interleukin signaling, macrophage function), “Cancer and Tumorigenesis” (e.g., molecular mechanisms of cancer, STAT3 signaling), “Growth and Development” (e.g., cardiac hypertrophy, GPCR signaling), and “Metabolic and Lipid Regulation” (e.g., PPAR signaling) showed divergent regulation between the two DEG sets. Ligand-independent pathways repressed these pathways, while ligand-dependent pathways generally showed activation of these pathways.

**Fig. 5.**
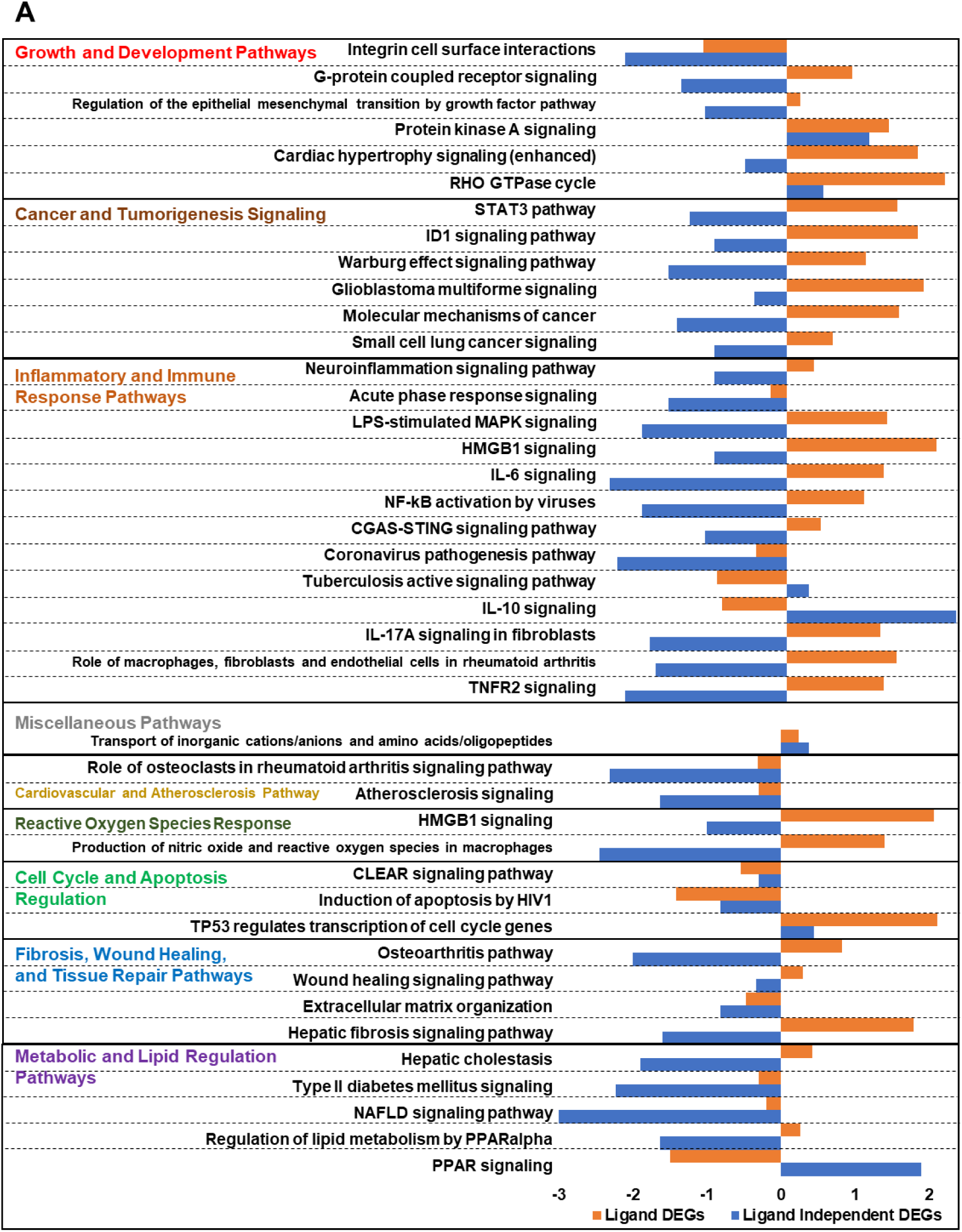
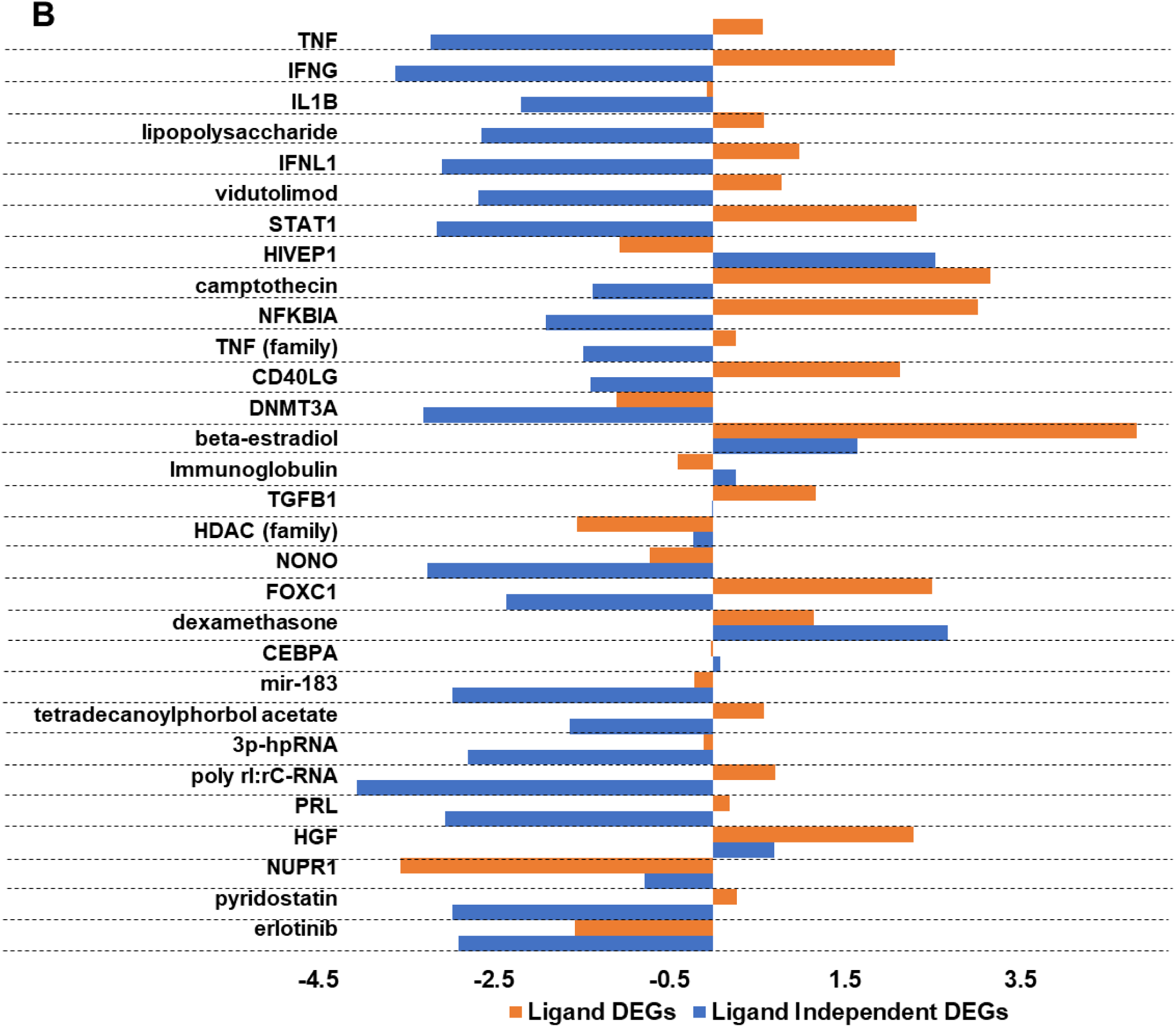
Divergent pathway and upstream regulator activity between ligand-receptor and ligand-independent VDR transcriptomes. (A) Comparison of canonical pathways enriched in 2,311 ligand-receptor DEGs and 599 ligand-independent DEGs. Of 43 overlapping pathways, only 13 showed consistent directionality (activation or inhibition), with the majority exhibiting divergent regulation in inflammation, cancer, growth, and metabolism-related pathways. (B) Number of predicted upstream regulators identified in ligand-receptor (3,300) and ligand-independent (604) DEG sets. Among 564 shared regulators, many (e.g., TNF, LPS, STAT1, FOXC1) showed opposing activation states, highlighting ligand-dependent modulation of key biological processes.

Analysis of the upstream regulators of these pathways shows a similar pattern (Fig. 5B). In the upstream regulator analysis, 3,300 regulators were predicted from the ligand-receptor DEGs, while 604 were predicted from the ligand-independent DEGs (Fig. 5B and Supplementary material table 3). The upstream regulators were impacted in the same direction by the ligand-independent and dependent pathways involved in hormone and growth factor signaling (dexamethasone, β-estradiol, and HGF) and epigenetic pathways (HDAC and DMNT3a). However, the majority of upstream regulators were shared between receptor and ligand DEGs, and their regulatory directions often differed. For instance, TNF, LPS, and IL1B were predicted as activators in receptor DEGs but as inhibitors in ligand DEGs (Fig. 5B). This divergence suggests that the ligand-independent action of the VDR is to repress these inflammatory pathways, while the ligand-dependent action is to initiate inflammatory pathways.

### 3.6. Cistromic modifications induced by 1,25(OH)_2_D_3_ in lentiviral transduced T-HESCs

We next investigated the cistromic changes induced by 1,25(OH)₂D₃ via CUT&RUN assays in T-HESCs. Cistromic analysis on the endogenous VDR in T-HESCs was unsuccessful due to the low levels of the receptor and the sensitivity of the antibody. To circumvent this, we transduced with pLenti-hVDR-HA, as described in the previous approach, to increase the sensitivity of the assay. pLenti-GFP was used as a control. Both transduced cells were treated with either vehicle or 1,25(OH)₂D₃. Peaks were normalized by subtracting signals from IgG controls and pLenti-GFP vector-transduced cells. Combined union peaks from technical triplicates were utilized for downstream analysis, excluding Chromosome Y peaks to eliminate false positives inherent to endometrial stromal cells. 1,25(OH)₂D₃ treatment significantly increased both the number and intensity of peaks compared to the vehicle, as demonstrated by heatmaps and metaplots (Fig. 6A). The number of normalized peaks for 1,25(OH)₂D₃-treated cells was 6,092, markedly higher than the vehicle’s 885 peaks (Fig. 6B). A total of 519 peaks were common between the two groups, accounting for approximately 58.64% of the vehicle peaks (Fig. 6B).

**Fig. 6.**
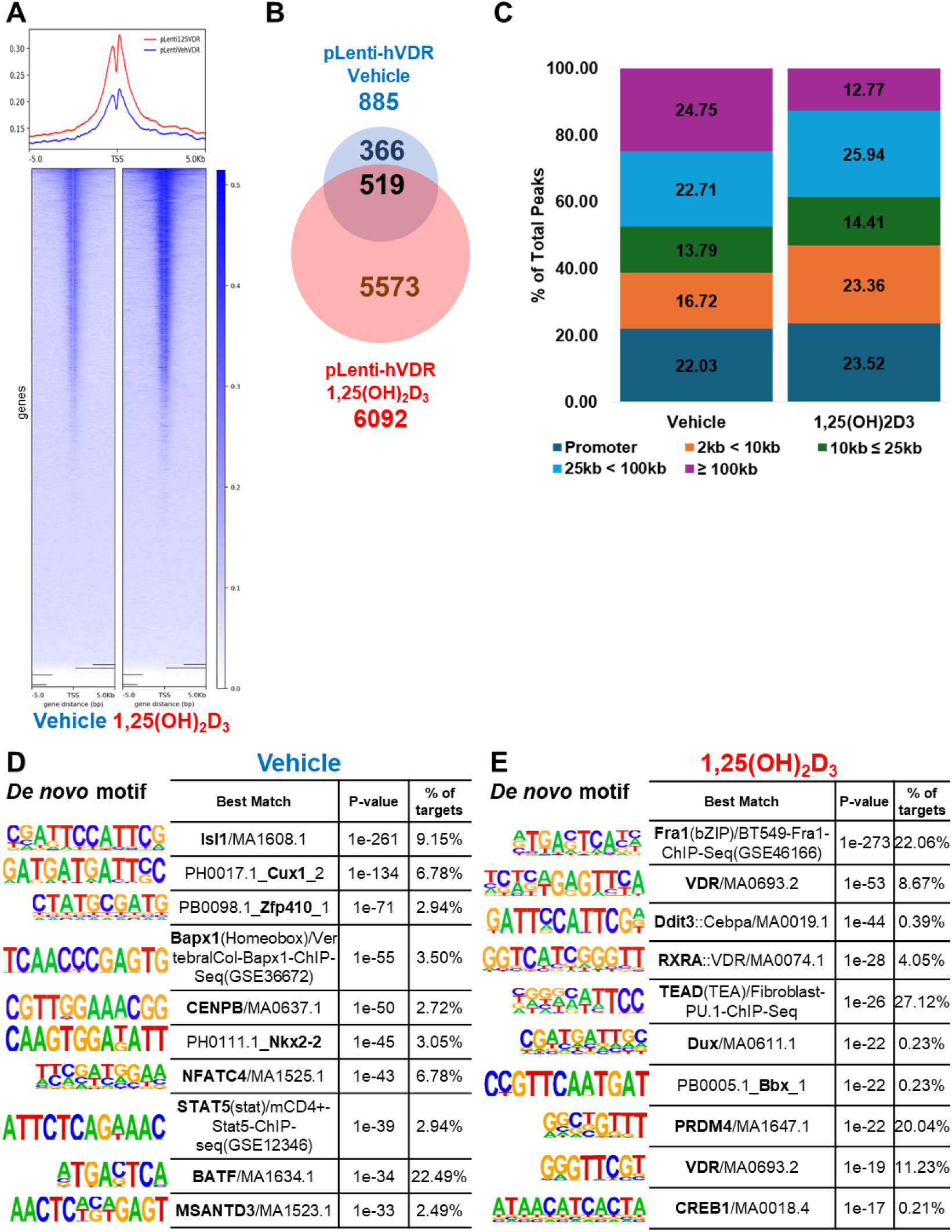
Cistromic modifications induced by 1,25(OH)₂D₃ in lentiviral activated T-HESCs. (A) Heatmap and metaplot of normalized CUT&RUN peaks showing increased peak intensity and (B) the number in 1,25(OH)₂D₃-treated cells compared to vehicle controls. (B) The number of overlapped peak locations between (C) Annotation of CUT&RUN peaks from vehicle- and 1,25(OH)₂D₃-treated cells. The charts illustrate the distribution of peaks across promoter, proximal, distal, and other genomic regions. HOMER motif analysis of CUT&RUN peaks from (D) vehicle- and (E) 1,25(OH)₂D₃-treated cells, highlighting *de novo* motifs and known enriched in each condition.

Peak annotation using MACS2 revealed distinct differences in the genomic distribution of CUT&RUN peaks between vehicle- and 1,25(OH)₂D₃-treated samples. Both conditions exhibited a substantial proportion of peaks localized to promoter regions, which were defined as being within 2 kb upstream of transcriptional start sites (TSS). Promoter peaks accounted for 22.03% of total peaks in vehicle-treated cells and 23.52% in 1,25(OH)₂D₃-treated cells (Fig. 6C and Supplementary material table 4). However, the distribution of peaks in distal regions, defined as ≥ ±100 kb from the TSS, showed a marked contrast. Distal peaks were significantly more frequent in vehicle-treated cells, representing 24.75% of the total peaks. In contrast, 1,25(OH)₂D₃-treated cells exhibited only 12.77% of their peaks in distal regions, indicating a relocation of binding sites away from distal genomic regions upon ligand activation (Fig. 6C and Supplementary material table 4).

The proportion of peaks in intergenic regions, typically non-coding areas between genes, also showed notable differences. In vehicle-treated cells, intergenic peaks constituted 41.36% of the total peaks (Supplementary material table 4). This proportion decreased to 30.22% in 1,25(OH)₂D₃-treated cells, further emphasizing a shift in chromatin binding toward gene-proximal areas (Supplementary material table 4). Peaks located in introns, exons, and other regulatory elements were less affected, suggesting that the ligand-induced redistribution predominantly impacts distal and intergenic regions. This shift in peak localization indicates that 1,25(OH)₂D₃ promotes the recruitment of VDR or other chromatin-associated proteins to regions near promoters and transcriptional start sites. Conversely, intronic, exonic, and proximal regulatory elements maintained relatively stable proportions (Supplementary material table 4). This redistribution suggests that 1,25(OH)₂D₃ facilitates the recruitment of VDR and associated chromatin factors toward promoters and transcriptional start sites, reducing suppressive binding in distal regions (Supplementary material table 4).

To further elucidate the specific DNA-binding motifs within these peaks, we conducted motif analysis using HOMER. In vehicle-treated cells, *de novo* motif analysis revealed significant enrichment for motifs associated with Isl1 (p-value 1e-261, 9.15% of target sites), followed by Cux1, Zfp410, Papx1, and CENPB. Known motif analysis identified Fra1, Atf3, Fra2, JunB, and Fosl2 as enriched (Fig. 6D and Supplementary material table 4). These motifs are consistent with a basal suppressive chromatin state commonly observed in the absence of ligand activation. In contrast, 1,25(OH)₂D₃-treated cells exhibited enrichment for motifs associated with Fra1, VDR, Ddit3, RXRA, and TEAD in *de novo* analysis (Fig. 6E and Supplementary material table 4). RXRA, the canonical heterodimer partner of VDR, was notably enriched, corroborating the ligand-activated recruitment of VDR-RXRA complexes to target regions. The presence of VDR and RXRA motifs exclusively in 1,25(OH)₂D₃-treated peaks underscores the ligand-dependent activation of VDR, which is otherwise ubiquitously present in endometrial stromal cells. Interestingly, known motif analysis for 1,25(OH)₂D₃-treated peaks still showed overlap with vehicle-enriched motifs, including Fra1, Fra2, Atf3, JunB, and Fos. These motifs are recognized for their roles in activating atrial natriuretic peptide gene transcription, suggesting that 1,25(OH)₂D₃ retains some basal chromatin features while facilitating selective VDR-driven transcriptional regulation (Supplementary material table 4). These findings underscore the dynamic nature of VDR-mediated chromatin remodeling in response to 1,25(OH)₂D₃. The observed redistribution of chromatin peaks from distal and intergenic regions to promoter-proximal sites, coupled with the selective enrichment of regulatory motifs, highlights VDR’s dual role as a basal suppressor and a ligand-activated transcriptional regulator in human endometrial stromal cells.

Since the Cut & Run analysis was conducted on virally expressed VDR, we compared these results to a previously published database on the VDR ChIP-Seq data set from the human kidney [66, 67]. From the kidney ChIP samples, we identified 30,113 unique peaks for subsequent analyses (Supplementary material table 5). Using the union of peaks from the kidney samples, we performed both *de novo* and known motif analyses with HOMER. ZNF354C emerged as the most significantly enriched binding motif, followed by Irf4, CHR, CEBPG, and VDR, and so on (Supplementary material figure 5A and Supplementary material table 5). Genomic annotation of the 30,113 peaks revealed that most were located in promoters (33.59%), and the distribution suggests that VDR transcription binds to sites similar to those it occupies when activated by 1,25(OH)₂D₃ exposure in T-HESCs (Supplementary material figure 5B). We overlapped the peaks from the T-HESC with the kidney-derived peaks. Among the vehicle-treated stromal cell peaks, 314 overlapped with kidney peaks (35.48%), while 1,25(OH)₂D₃-treated stromal cell peaks showed a 32.27% overlap (1,966 peaks) (Supplementary material figure 5C). These results suggest that VDR binds similar genomic regions in both its basal and ligand-bound states and confirm the reliability of VDR binding in endometrial stromal cells.

### 3.7. Epigenetic atlas of chromatin, transcriptomic, and cistromic networks mediating VDR function in human endometrial stromal cells

To investigate the interplay between cistromic binding patterns and transcriptomic changes, we previously analyzed VDR-binding peaks identified *via* CUT&RUN in pLenti-hVDR cells treated with either vehicle or 1,25(OH)₂D₃ (Fig. 6A). Based on the previous cistromic data, we annotated the nearest genes from each peak region. Among ligand-independent DEGs (599 genes), only 39 genes overlapped with basal VDR-binding peaks, representing 6.51% (Fig. 7A). In contrast, ligand-treated DEGs (2,311 genes; siNT 1,25(OH)₂D₃ vs. siVDR 1,25(OH)₂D₃) demonstrated a significantly higher overlap, with 1,052 genes (45.52%) intersecting with 1,25(OH)₂D₃-associated peaks (Fig. 7B). These findings suggest that, in the absence of the ligand, VDR may bind DNA but does not robustly regulate transcription. This lack of overlap may reflect the recruitment of VDR to distal regulatory elements, such as enhancers or silencers, which remain transcriptionally inert without ligand activation.

**Fig. 7.**
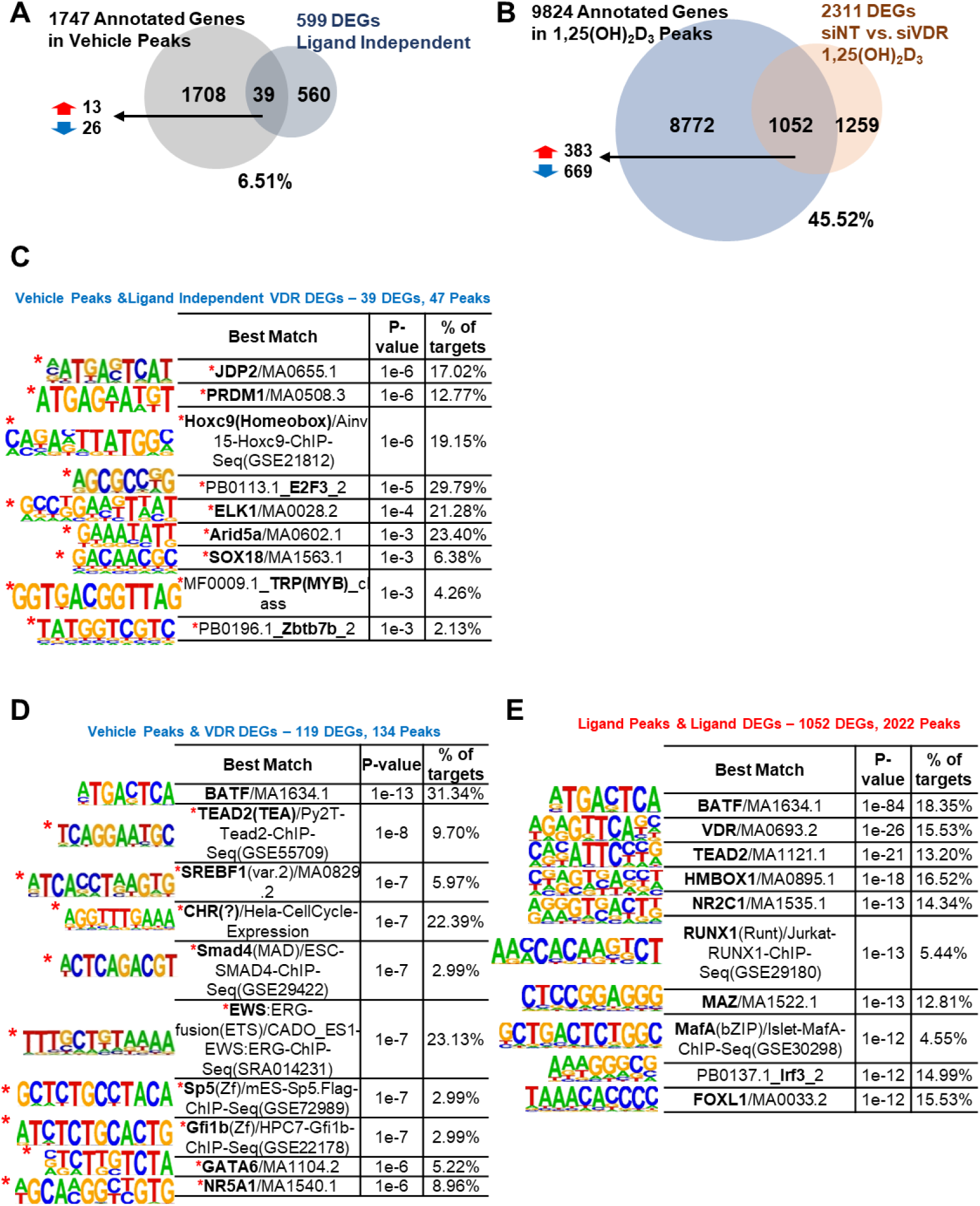
Interplay between cistromic peaks and transcriptomic changes in VDR-mediated regulation of T-HESCs. Vehicle-treated cistromic peaks annotated 1,747 genes. (A) VDR cistromic peaks with ligand-independent receptor DEGs representing 39 (6.51%) genes. (B) Annotated genes from the ligand peaks with ligand-receptor DEGs (siNT 1,25(OH)₂D₃ vs. siVDR 1,25(OH)₂D₃). 9,824 genes (45.52%) annotated from 1,25(OH)₂D₃ peaks overlapped with ligand-receptor DEGs. HOMER motif analyses of VDR peaks within (C) ligand-independent receptor DEGs, (D) receptor DEGs (vehicle-treated), and (E) VDR peaks within ligand DEGs (1,25(OH)₂D₃-treated).

### 3.8. Functional significance of VDR cistromic activity in DEGs

To dissect the localization of VDR-binding peaks within DEG-associated loci, we performed genomic annotation analyses. Basal peaks within receptor DEGs were distributed as follows: promoters (26.87%), 25–100 kb upstream (28.36%), <10 kb upstream (22.39%), 10–25 kb upstream (15.67%), and ≥100 kb upstream (6.72%) (Supplementary material table 6). Similarly, ligand-induced peaks within ligand DEGs were located in promoters (25.42%), 25–100 kb upstream (27.10%), <10 kb upstream (25.32%), 10–25 kb upstream (16.02%), and ≥100 kb upstream (6.13%) (Supplementary material table 6). These consistent distributions suggest that VDR regulates both proximal and distal regulatory regions.

Motif analysis of VDR peaks in DEGs revealed functional distinctions. For ligand-independent receptor DEGs (39 genes), 47 peaks revealed motifs for JDP2, PROM1, HOXC9, E2F3, ELK1, ARID5A, SOX18, TRP, and ZBTB7B (Fig. 7C). These genes participate in proliferation (ID3, HMGA1, ATF3), ECM remodeling (SPP1, FAT4), and immune responses (TNFRSF6B, GDNF), highlighting their roles in tissue homeostasis and pathophysiology. For broader receptor DEGs, 119 genes yielded 134 peaks with motifs for BATF, TEAD2, SREBF1, CHR, SMAD4, EWS, SP5, GFI1B, GATA6, and NR5A1 (Fig. 7D), suggesting basal regulation of stromal identity to maintain tissue homeostasis responding to injury and potentially contributing to pathological conditions.

For ligand-treated cells, motifs for BATF, VDR, TEAD2, HMBOX1, NR2C1, RUNX1, MAZ, MAFA, IRF3, and FOXL1 were highly enriched, implicating roles in decidualization, immune modulation, and structural organization (Fig. 7E and Supplementary material table 6). These results suggest that ligand-activated VDR operates as part of a coordinated transcription factor network, enabling context-dependent gene regulation.

## 4. Discussion

This study provides novel insights into the role of active vitamin D (1,25(OH)_2_D_3_) and its receptor (VDR) in uterine biology, offering evidence that vitamin D deficiency impairs uterine function and that VDR regulates endometrial stromal cell differentiation. To our knowledge, this is the first report of a correlation between a vitamin D-deficient diet and impaired decidualization in an in vivo mouse model, together with evidence suggesting a potential suppressor role for VDR in human endometrial stromal cell differentiation. These findings represent a significant advance in understanding the molecular and epigenetic mechanisms governing endometrial health and their implications for female reproductive health.

Our study demonstrates that vitamin D deficiency severely impairs uterine decidualization, a key process for successful pregnancy. Mice maintained on a vitamin D-deficient diet exhibited diminished uterine weight ratios during decidualization and downregulation of pivotal decidual markers, including Wnt4 [68–70], which is essential for endometrial receptivity. This is the first in vivo evidence directly linking dietary vitamin D deficiency to impaired decidualization, emphasizing the unique role of vitamin D in modulating uterine function in mice. In addition to biochemical and molecular alterations, vitamin D-deficient dams experienced heightened reproductive challenges, including increased labor difficulties and maternal mortality. Collectively, these results indicate that vitamin D deficiency may cause adverse fetal and maternal outcomes by impairing decidualization.

In human endometrial stromal cells (T-HESCs), our data further support a regulatory role for VDR in decidualization. We observed that VDR expression decreased during hormone-induced decidualization, suggesting its suppression may be necessary for effective differentiation. Functional assays revealed that VDR acts as a suppressor of differentiation: silencing VDR enhanced decidual marker expression, while VDR overexpression diminished it. These findings significantly contribute to understanding VDR as a context-dependent regulator, particularly in its suppressive role during early stromal differentiation. This is the first report implicating VDR as a suppressor in human endometrial stromal cell differentiation. It raises critical questions about whether and how VDR fine-tunes the transition of the endometrium from a proliferative to a differentiated state. The results also suggest a potential balance between the suppressive and supportive roles of VDR, contingent on its ligand, 1,25(OH)₂D₃. Future studies will be required to unravel this dynamic and explore its implications for decidualization and other reproductive processes.

Chromatin accessibility assays revealed that VDR influences the epigenomic landscape by suppressing transcriptionally active chromatin states. Specifically, in the absence of ligand activation, VDR knockdown resulted in heightened chromatin accessibility (Fig. 3A and B) and increased deposition of active histone marks, including H3K4me3 [71] and H3K27ac [72] (Fig. 3C). This suggests that VDR maintains a transcriptionally repressive chromatin state under basal conditions, supporting its role as a suppressor. Importantly, ligand activation with 1,25(OH)₂D₃ reprogrammed VDR binding patterns, increasing receptor occupancy at proximal sites (Fig. 6C). This redistribution facilitated transcriptional activation of differentiation-associated genes, highlighting the dynamic role of VDR in chromatin remodeling. The interplay between these chromatin-level changes and transcriptional regulation underscores the epigenetic versatility of VDR in response to vitamin D signaling. To further delineate the regulatory network underlying VDR function, we performed motif enrichment analysis on VDR-bound regions overlapping VDR-dependent DEGs (all without ligand). This analysis revealed several significantly enriched transcription factor binding motifs. Among the top hits was BATF, a transcription factor known to coordinate chromatin remodeling during immune cell differentiation [73, 74]. Its presence in over 30% of target regions suggests BATF may act in concert with VDR to modulate chromatin accessibility, potentially by recruiting chromatin-modifying complexes. Together, these findings indicate that VDR modulates gene expression by influencing or cooperating with other transcription factors that regulate chromatin accessibility.

Analysis of the transcriptomic impact of VDR and its ligand on endometrial stromal cell biology demonstrates that ligand-independent effects of VDR repress inflammatory pathways, whereas treatment with the ligand activates them. Inflammation plays a crucial role in decidualization and is necessary for stromal differentiation [75, 76]. The role of VDR may be to maintain endometrial cells in an undifferentiated state until the appropriate stimulus initiates decidualization. Thus, VDR and its ligand act as a gatekeeper for endometrial stromal biology, protecting against premature differentiation while facilitating differentiation when appropriate stimuli are present.

There are 27 chromatin-related receptor DEGs (Supplementary material figure 6A). GSEA analysis revealed strong associations with Gene Ontology (GO) biological processes such as regulation of chromosome organization, cell cycle checkpoint signaling, sister chromatid segregation, and nuclear chromosome segregation (Supplementary material figure 6B and C). Specifically, these chromatin-related GO terms showed highly suppressed normalized enrichment scores in chromatin organization and remodeling (Supplementary material figure 6D).

Our cistromic analysis identified canonical VDR-RXRA motifs in VDR-binding regions [77–79], corroborating its established heterodimeric interaction with RXR (Fig. 6E). Ligand activation significantly altered VDR-binding patterns, enhancing accessibility at proximal regions of key decidualization genes. To our knowledge, this is the first study integrating chromatin accessibility, histone modifications, and cistromic analysis to dissect VDR’s role in human endometrial stromal cells. Paradoxically, VDR knockdown in ligand-treated cells amplified chromatin accessibility, even at regions where VDR typically suppresses transcription. This suggests a dual function for VDR as both a repressor and activator, with its role dictated by ligand presence and context (Fig. 6D and E). These findings provide new perspectives on VDR’s role as a chromatin-level modulator and its broader implications for reproductive biology.

Our findings reveal the dual role of vitamin D in female reproduction. While basal VDR restrains decidualization by controlling chromatin accessibility and repressing inflammation, its activation by vitamin D promotes differentiation. These results suggest that vitamin D supplementation could be a therapeutic strategy for improving uterine function and pregnancy outcomes in women with vitamin D deficiency [80, 81]. This may be particularly relevant given the low levels of 25(OH)D in U.S. women of reproductive age [82]. The suppressive role of VDR in endometrial stromal cell differentiation also raises intriguing possibilities for its involvement in other processes, such as implantation failure or susceptibility to endometrial disorders like endometriosis. Moreover, the chromatin remodeling effects mediated by VDR highlight its potential influence on the epigenomic landscape of the endometrium. These findings provide a foundation for future research into vitamin D and VDR as key modulators of endometrial plasticity and health.

Future research should focus on elucidating the downstream targets of VDR-mediated transcriptional regulation during decidualization. A combination of transcriptomic, proteomic, and metabolomic analyses will likely yield deeper insights into VDR’s broader impact on endometrial physiology. Additionally, potential interactions between VDR and other signaling pathways, such as estrogen and progesterone receptors, warrant further exploration to clarify their cooperative roles in endometrial biology [83, 84]. Clinical studies should also investigate the efficacy of vitamin D supplementation in enhancing decidualization and improving pregnancy outcomes in women with vitamin D deficiency. Such trials should consider genetic and epigenetic variations in VDR that may influence individual responses to supplementation [85]. Furthermore, extending these findings to other reproductive tissues, such as the ovary and placenta, could uncover novel therapeutic targets for reproductive disorders.

## 5. Conclusion

This study reveals a dual, ligand-dependent role of VDR in female reproduction. Vitamin D deficiency impaired decidualization in vivo. In its unliganded state, VDR suppresses differentiation in human endometrial stromal cells by modulating chromatin accessibility and repressing inflammation. Upon activation by 1,25(OH)_2_D_3_, however, VDR promotes this differentiation process by activating inflammation. By integrating chromatin-level analyses with functional studies, we have uncovered novel mechanisms by which vitamin D and VDR regulate uterine function. These insights lay the groundwork for future research into vitamin D’s broader implications in reproductive health and beyond.

## CRediT authorship contribution statement

**MyeongJin Yi:** Conceptualization, Data curation, Formal analysis, Investigation, Methodology, Validation, Visualization, Writing - original draft, Writing – review & editing. **Skylar G. Montague Redecke:** Formal analysis, Writing – review & editing. **Tianyuan Wang:** Data curation, Formal analysis, Software, Validation. **Austin Bell-Hensley:** Data curation, Formal analysis, Software, Validation. **Shuyun Li:** Formal analysis, Writing – review & editing. **Abdull J. Massri:** Data curation, Formal analysis, Software. **Anne Marie Z. Jukic:** Conceptualization, Funding acquisition, Writing – review & editing. **Francesco J. DeMayo:** Conceptualization, Funding acquisition, Resources, Supervision, Writing-review & editing.

## Acknowledgments

This research was supported in part by the Intramural Research Program of the National Institutes of Health (NIH). The contributions of the NIH author(s) were made as part of their official duties as NIH federal employees, are in compliance with agency policy requirements, and are considered Works of the United States Government. However, the findings and conclusions presented in this paper are those of the author(s) and do not necessarily reflect the views of the NIH or the U.S. Department of Health and Human Services.

The authors appreciate the support from the Veterinary Medicine Section (ZIGES102825), Viral Vector Core (ZICES102506), Epigenomics and DNA Sequencing Core (ZICES102545), Protein Expression Facility (ZICES102487), and Integrative Bioinformatics Supportive Group (ZICES103371) at the National Institute of Environmental Health Sciences (NIEHS). Also, technical advice for CUT&RUN analysis was provided by Dr. Krystal Orlando at the Epigenetics and RNA Biology Laboratory (ERBL) at NIEHS.

## Appendix A Supplementary material

Supplementary methods and figures

## Appendix B Supplementary spreadsheets

Supplementary Table 1. Key sources, information on the nutrition of vitamin D diets, and a list of abbreviations.

Supplementary Table 2. Information on peaks within open chromatin regions in T-HESCs, both siNT and siVDR.

Supplementary Table 3. List of DEGs, canonical pathways, and upstream regulators in T-HESCs regulated by ligand (1,25(OH)_2_D_3_) and receptor (VDR).

Supplementary Table 4. Information on peaks and motifs in lentiviral transduced T-HESCs treated with either vehicle or 1,25(OH)_2_D_3_.

Supplementary Table 5. Information on peaks and motifs in the human kidney (GSE129585) and overlay peaks and motifs between human kidney and lentiviral transduced T-HESCs treated with either vehicle or 1,25(OH)_2_D_3_.

Supplementary Table 6. List of overlapped peaks, DEGs, and motifs between cistromic and transcriptomic analyses.

## Supplementary data statement

Supplementary data associated with this article can be found in the online version.

## Data availability

The raw and processed data files generated in this study are available in NCBI GEO with the accession numbers: RNA-seq (GSE254251), ATAC-seq (GSE306094), and Cut&Run-seq (GSE306127). The following datasets were generated:

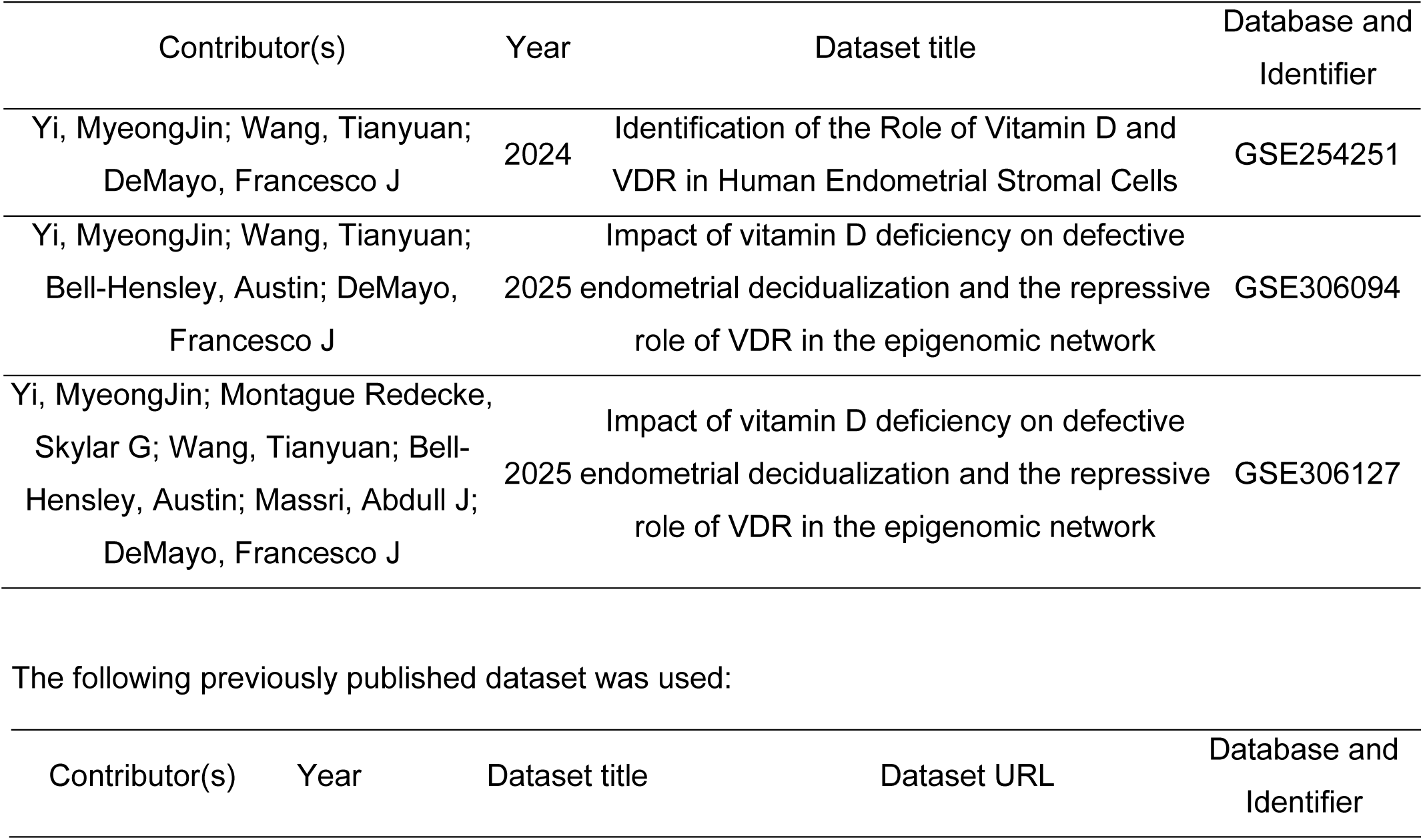
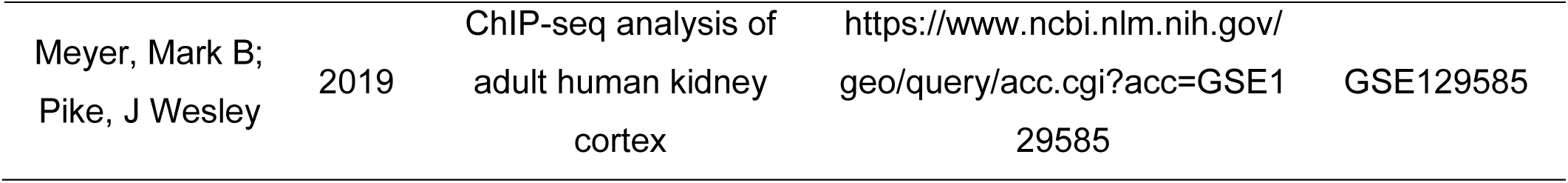

| Contributor(s) | Year | Dataset title | Database and Identifier |
| --- | --- | --- | --- |
| Yi, MyeongJin; Wang, Tianyuan; DeMayo, Francesco J | 2024 | Identification of the Role of Vitamin D and VDR in Human Endometrial Stromal Cells | GSE254251 |
| Yi, MyeongJin; Wang, Tianyuan; Bell-Hensley, Austin; DeMayo, Francesco J | 2025 | Impact of vitamin D deficiency on defective endometrial decidualization and the repressive role of VDR in the epigenomic network | GSE306094 |
| Yi, MyeongJin; Montague Redecke, Skylar G; Wang, Tianyuan; Bell-Hensley, Austin; Massri, Abdull J; DeMayo, Francesco J | 2025 | Impact of vitamin D deficiency on defective endometrial decidualization and the repressive role of VDR in the epigenomic network | GSE306127 |

The following previously published dataset was used:
| Contributor(s) | Year | Dataset title | Dataset URL | Database and Identifier |
| --- | --- | --- | --- | --- |

|  |  |  |  |  |
| --- | --- | --- | --- | --- |
| Meyer, Mark B;<br>Pike, J Wesley | 2019 | ChIP-seq analysis of<br>adult human kidney<br>cortex | https://www.ncbi.nlm.nih.gov/<br>geo/query/acc.cgi?acc=GSE1<br>29585 | GSE129585 |

## References

[1] L. Shi, Y. Bao, X. Deng, X. Xu, J. Hu, Association between calcium and vitamin D supplementation and increased risk of kidney stone formation in patients with osteoporosis in Southwest China: a cross-sectional study, BMJ Open 15(2) (2025) e092901.

[2] T. Al-Barazenji, A. Allouch, N. Al Husaini, S. Yousef, W.N. Ibrahim, A. Al-Haidose, H. Zayed, A.M. Abdallah, Association Between Vitamin D Receptor BsmI Polymorphism and Low Bone Mineral Density in Postmenopausal Women in the MENA Region, Pathophysiology 32(1) (2025).

[3] A.M.Z. Jukic, Q.E. Harmon, Accumulating evidence for vitamin D and conception, Fertil Steril 113(2) (2020) 330–331.

[4] X. Cui, R.A.N. Pertile, V. Raman, D. Eyles, Vitamin D differentiates dopamine neurons in vitro, increasing neurite architecture, dopamine release and expression of relevant synaptic proteins, J Steroid Biochem Mol Biol 247 (2025) 106681.

[5] H. Evans, A. Greenhough, L. Perry, G. Lasanta, C.M. Gonzalez, A. Mourino, J.P. Mansell, Hypoxia Compromises the Differentiation of Human Osteosarcoma Cells to CAR-R, a Hydroxylated Derivative of Lithocholic Acid and Potent Agonist of the Vitamin D Receptor, Int J Mol Sci 26(1) (2025).

[6] C.M. Wang, Y.J. Chen, B.C. Yang, J.W. Yang, W. Wang, Y. Zeng, J. Jiang, Supplementation with active vitamin D3 ameliorates experimental autoimmune thyroiditis in mice by modulating the differentiation and functionality of intrathyroidal T-cell subsets, Front Immunol 16 (2025) 1528707.

[7] M. Zhang, F. Zhao, M. Guo, M. Duan, Y. Xie, L. Qiu, Vitamin E alleviates zebrafish intestinal damage and microbial disturbances caused by pyraclostrobin, Pestic Biochem Physiol 208 (2025) 106221.

[8] M.S. Sater, Z.H.A. Malalla, M.E. Ali, H.A. Giha, Downstream Link of Vitamin D Pathway with Inflammation Irrespective of Plasma 25OHD3: Hints from Vitamin D-Binding Protein (DBP) and Receptor (VDR) Gene Polymorphisms, Biomedicines 13(2) (2025).

[9] P. Skubica, I. Hoffmanova, P. Dankova, Chronically increased osteoclastogenesis in adult celiac disease patients does not hinder improvement in bone health induced by gluten-free diet: Role of vitamin D, OPG and IL-6, J Nutr Biochem (2025) 109871.

[10] R. Heidarzadehpilehrood, H.A. Hamid, M. Pirhoushiaran, Vitamin D receptor (VDR) gene polymorphisms and risk for polycystic ovary syndrome and infertility: An updated systematic review and meta-analysis, Metabol Open 25 (2025) 100343.

[11] X. Han, H. Yang, Evaluation of placental growth factor, Vitamin D, and systemic inflammatory index as predictive biomarkers for preeclampsia severity: a retrospective cohort study, BMC Pregnancy Childbirth 25(1) (2025) 75.

[12] S.L. Mumford, R.A. Garbose, K. Kim, K. Kissell, D.L. Kuhr, U.R. Omosigho, N.J. Perkins, N. Galai, R.M. Silver, L.A. Sjaarda, T.C. Plowden, E.F. Schisterman, Association of preconception serum 25-hydroxyvitamin D concentrations with livebirth and pregnancy loss: a prospective cohort study, Lancet Diabetes Endocrinol 6(9) (2018) 725–732.

[13] A.M.Z. Jukic, D.D. Baird, C.R. Weinberg, A.J. Wilcox, D.R. McConnaughey, A.Z. Steiner, Pre-conception 25-hydroxyvitamin D (25(OH)D) and fecundability, Hum Reprod 34(11) (2019) 2163–2172.

[14] H. Hosseinirad, S. Paktinat, F. Mohanazadeh Falahieh, M. Mirani, A. Karamian, A. Karamian, Z. Shams Mofarahe, Effect of 1,25(OH)2-vitamin D3 on decidualization of human endometrial stromal cells, Steroids 180 (2022) 108978.

[15] H. Du, G.S. Daftary, S.I. Lalwani, H.S. Taylor, Direct regulation of HOXA10 by 1,25-(OH)2D3 in human myelomonocytic cells and human endometrial stromal cells, Mol Endocrinol 19(9) (2005) 2222–33.

[16] S.Y. Chan, R. Susarla, D. Canovas, E. Vasilopoulou, O. Ohizua, C.J. McCabe, M. Hewison, M.D. Kilby, Vitamin D promotes human extravillous trophoblast invasion in vitro, Placenta 36(4) (2015) 403–9.

[17] W. Yang, Z. Hu, W. Gu, Assessing the relationship between serum vitamin A, C, E, D, and B12 levels and preeclampsia, J Matern Fetal Neonatal Med 38(1) (2025) 2466222.

[18] W.C. Yang, R. Chitale, K.M. O’Callaghan, C.R. Sudfeld, E.R. Smith, The Effects of Vitamin D Supplementation During Pregnancy on Maternal, Neonatal, and Infant Health: A Systematic Review and Meta-analysis, Nutr Rev 83(3) (2025) e892–e903.

[19] C. Lin, H. Liu, C. Chen, B. Fu, Correlation between HOMA-IR and Pregnancy Outcomes of GDM Patients Under Vitamin D Insufficiency or Deficiency, Clin Lab 71(2) (2025).

[20] T. Zhang, L. Yang, S. Yang, S. Gao, Vitamin D on the susceptibility of gestational diabetes mellitus: a mini-review, Front Nutr 12 (2025) 1514148.

[21] C. Lin, H. Liu, Corrigendum: Relationship between vitamin D deficiency and gestational diabetes: a narrative review, Front Endocrinol (Lausanne) 15 (2024) 1548353.

[22] A.M.Z. Jukic, A.J. Wilcox, D.R. McConnaughey, C.R. Weinberg, A.Z. Steiner, 25-Hydroxyvitamin D and Long Menstrual Cycles in a Prospective Cohort Study, Epidemiology 29(3) (2018) 388–396.

[23] Q.E. Harmon, K. Kissell, A.M.Z. Jukic, K. Kim, L. Sjaarda, N.J. Perkins, D.M. Umbach, E.F. Schisterman, D.D. Baird, S.L. Mumford, Vitamin D and Reproductive Hormones Across the Menstrual Cycle, Hum Reprod 35(2) (2020) 413–423.

[24] L. Pal, H. Zhang, J. Williams, N.F. Santoro, M.P. Diamond, W.D. Schlaff, C. Coutifaris, S.A. Carson, M.P. Steinkampf, B.R. Carr, P.G. McGovern, N.A. Cataldo, G.G. Gosman, J.E. Nestler, E. Myers, R.S. Legro, N. Reproductive Medicine, Vitamin D Status Relates to Reproductive Outcome in Women With Polycystic Ovary Syndrome: Secondary Analysis of a Multicenter Randomized Controlled Trial, J Clin Endocrinol Metab 101(8) (2016) 3027–35.

[25] M. Seyyed Abootorabi, P. Ayremlou, T. Behroozi-Lak, S. Nourisaeidlou, The effect of vitamin D supplementation on insulin resistance, visceral fat and adiponectin in vitamin D deficient women with polycystic ovary syndrome: a randomized placebo-controlled trial, Gynecol Endocrinol 34(6) (2018) 489–494.

[26] B. Zhang, X. Yao, X. Zhong, Y. Hu, J. Xu, Vitamin D supplementation in the treatment of polycystic ovary syndrome: A meta-analysis of randomized controlled trials, Heliyon 9(3) (2023) e14291.

[27] M. Yang, X. Shen, D. Lu, J. Peng, S. Zhou, L. Xu, J. Zhang, Effects of vitamin D supplementation on ovulation and pregnancy in women with polycystic ovary syndrome: a systematic review and meta-analysis, Front Endocrinol (Lausanne) 14 (2023) 1148556.

[28] P. Deryabin, A. Griukova, N. Nikolsky, A. Borodkina, The link between endometrial stromal cell senescence and decidualization in female fertility: the art of balance, Cell Mol Life Sci 77(7) (2020) 1357–1370.

[29] H. Gong, W. Wu, J. Xu, D. Yu, B. Qiao, H. Liu, B. Yang, Y. Li, Y. Ling, H. Kuang, Flutamide ameliorates uterine decidualization and angiogenesis in the mouse hyperandrogenemia model during mid-pregnancy, PLoS One 14(5) (2019) e0217095.

[30] J. Filant, T.E. Spencer, Uterine glands: biological roles in conceptus implantation, uterine receptivity and decidualization, Int J Dev Biol 58(2-4) (2014) 107–16.

[31] W. Zhao, Y. Wang, J. Liu, Q. Yang, S. Zhang, X. Hu, Z. Shi, Z. Zhang, J. Tian, D. Chu, L. An, Progesterone Activates the Histone Lactylation-Hif1alpha-glycolysis Feedback Loop to Promote Decidualization, Endocrinology 165(1) (2023).

[32] S. Chen, A. Zhang, N. Li, H. Wu, Y. Li, S. Liu, Q. Yan, Elevated high-mannose N-glycans hamper endometrial decidualization, iScience 26(11) (2023) 108170.

[33] H. Zhao, N. Lv, J. Cong, G. Chen, H. Bao, X. Liu, Upregulated RPA2 in endometrial tissues of repeated implantation failure patients impairs the endometrial decidualization, J Assist Reprod Genet 40(11) (2023) 2739–2750.

[34] M.A. Ochoa-Bernal, A.T. Fazleabas, Physiologic Events of Embryo Implantation and Decidualization in Human and Non-Human Primates, Int J Mol Sci 21(6) (2020).

[35] R. Thapa, L. Druessel, L. Ma, D.S. Torry, B.M. Bany, ATOH8 Expression Is Regulated by BMP2 and Plays a Key Role in Human Endometrial Stromal Cell Decidualization, Endocrinology 165(1) (2023).

[36] M.E. Diessler, R. Hernandez, G. Gomez Castro, C.G. Barbeito, Decidual cells and decidualization in the carnivoran endotheliochorial placenta, Front Cell Dev Biol 11 (2023) 1134874.

[37] C. Dunk, M. Kwan, A. Hazan, S. Walker, J.K. Wright, L.K. Harris, R.L. Jones, S. Keating, J.C.P. Kingdom, W. Whittle, C. Maxwell, S.J. Lye, Failure of Decidualization and Maternal Immune Tolerance Underlies Uterovascular Resistance in Intra Uterine Growth Restriction, Front Endocrinol (Lausanne) 10 (2019) 160.

[38] H. Okada, T. Tsuzuki, H. Murata, Decidualization of the human endometrium, Reprod Med Biol 17(3) (2018) 220–227.

[39] H.J. Kliman, Uteroplacental blood flow. The story of decidualization, menstruation, and trophoblast invasion, Am J Pathol 157(6) (2000) 1759–68.

[40] T. Yoshizawa, Y. Handa, Y. Uematsu, S. Takeda, K. Sekine, Y. Yoshihara, T. Kawakami, K. Arioka, H. Sato, Y. Uchiyama, S. Masushige, A. Fukamizu, T. Matsumoto, S. Kato, Mice lacking the vitamin D receptor exhibit impaired bone formation, uterine hypoplasia and growth retardation after weaning, Nat Genet 16(4) (1997) 391–6.

[41] N. Rashidi, S. Arefi, M. Sadri, A.A. Delbandi, Effect of active vitamin D on proliferation, cell cycle and apoptosis in endometriotic stromal cells, Reprod Biomed Online 46(3) (2023) 436–445.

[42] J.L. Kelsey, Methods in observational epidemiology, 2nd ed., Oxford University Press, New York, 1996.

[43] E. Quiroz, R.M. Marquardt, S.Y. Li, A. Gruzdev, D. Cunefare, C. Ganta, S.P. Wu, J.P. Lydon, F.J. DeMayo, A mouse model engineered to spatiotemporally control Cre expression in progesterone receptor positive cellsdagger, Biol Reprod 113(1) (2025) 83–96.

[44] M.J. Dickson, Y. Oh, A. Gruzdev, R. Li, N. Balaguer, A.M. Kelleher, T.E. Spencer, S.P. Wu, F.J. DeMayo, Inserting Cre recombinase into the Prolactin 8a2 gene for decidua-specific recombination in mice, Genesis 60(4-5) (2022) e23473.

[45] V.K. Maurya, M.M. Szwarc, D.M. Lonard, W.E. Gibbons, S.P. Wu, B.W. O’Malley, F.J. DeMayo, J.P. Lydon, Decidualization of human endometrial stromal cells requires steroid receptor coactivator-3, Front Reprod Health 4 (2022) 1033581.

[46] S.A. Michalski, S.B. Chadchan, E.S. Jungheim, R. Kommagani, Isolation of Human Endometrial Stromal Cells for In Vitro Decidualization, J Vis Exp (139) (2018).

[47] R. Li, T.Y. Wang, E. Shelp-Peck, S.P. Wu, F.J. DeMayo, The single-cell atlas of cultured human endometrial stromal cells, F S Sci 3(4) (2022) 349–366.

[48] M.M. Szwarc, L. Hai, W.E. Gibbons, M.C. Peavey, L.D. White, Q. Mo, D.M. Lonard, R. Kommagani, R.B. Lanz, F.J. DeMayo, J.P. Lydon, Human endometrial stromal cell decidualization requires transcriptional reprogramming by PLZF, Biol Reprod 98(1) (2018) 15–27.

[49] M.M. Szwarc, L. Hai, W.E. Gibbons, Q. Mo, R.B. Lanz, F.J. DeMayo, J.P. Lydon, Early growth response 1 transcriptionally primes the human endometrial stromal cell for decidualization, J Steroid Biochem Mol Biol 189 (2019) 283–290.

[50] J.J. Brosens, N. Hayashi, J.O. White, Progesterone receptor regulates decidual prolactin expression in differentiating human endometrial stromal cells, Endocrinology 140(10) (1999) 4809–20.

[51] D.J. Fowler, K.H. Nicolaides, J.P. Miell, Insulin-like growth factor binding protein-1 (IGFBP-1): a multifunctional role in the human female reproductive tract, Hum Reprod Update 6(5) (2000) 495–504.

[52] M. Harada, Y. Osuga, Y. Takemura, O. Yoshino, K. Koga, Y. Hirota, T. Hirata, C. Morimoto, T. Yano, Y. Taketani, Mechanical stretch upregulates IGFBP-1 secretion from decidualized endometrial stromal cells, Am J Physiol Endocrinol Metab 290(2) (2006) E268–72.

[53] J.R. Huang, L. Tseng, P. Bischof, O.A. Janne, Regulation of prolactin production by progestin, estrogen, and relaxin in human endometrial stromal cells, Endocrinology 121(6) (1987) 2011–7.

[54] J.D. Buenrostro, B. Wu, H.Y. Chang, W.J. Greenleaf, ATAC-seq: A Method for Assaying Chromatin Accessibility Genome-Wide, Curr Protoc Mol Biol 109 (2015) 21 29 1–21 29 9.

[55] J.D. Buenrostro, P.G. Giresi, L.C. Zaba, H.Y. Chang, W.J. Greenleaf, Transposition of native chromatin for fast and sensitive epigenomic profiling of open chromatin, DNA-binding proteins and nucleosome position, Nat Methods 10(12) (2013) 1213–8.

[56] T. Liu, Use model-based Analysis of ChIP-Seq (MACS) to analyze short reads generated by sequencing protein-DNA interactions in embryonic stem cells, Methods Mol Biol 1150 (2014) 81–95.

[57] Y. Zhang, T. Liu, C.A. Meyer, J. Eeckhoute, D.S. Johnson, B.E. Bernstein, C. Nusbaum, R.M. Myers, M. Brown, W. Li, X.S. Liu, Model-based analysis of ChIP-Seq (MACS), Genome Biol 9(9) (2008) R137.

[58] S.G. Montague Redecke, A. Bell-Hensley, S. Li, M. Yi, A. Jain, A.J. Massri, F.J. DeMayo, Deciphering estrogen receptor alpha-driven transcription in human endometrial stromal cells via transcriptome, cistrome, and integration with chromatin landscape, F S Sci (2025).

[59] S.G. Montague Redecke, A. Bell-Hensley, M. Yi, S. Li, F.J. DeMayo, PGR Isoform modulation via enhancer activation regulates progesterone signaling in endometrial stromal cells, F S Sci (2025).

[60] R. Li, T. Wang, R.M. Marquardt, J.P. Lydon, S.P. Wu, F.J. DeMayo, TRIM28 modulates nuclear receptor signaling to regulate uterine function, Nat Commun 14(1) (2023) 4605.

[61] T. Downey, Analysis of a multifactor microarray study using Partek genomics solution, Methods Enzymol 411 (2006) 256–70.

[62] S. Heinz, C. Benner, N. Spann, E. Bertolino, Y.C. Lin, P. Laslo, J.X. Cheng, C. Murre, H. Singh, C.K. Glass, Simple combinations of lineage-determining transcription factors prime cis-regulatory elements required for macrophage and B cell identities, Mol Cell 38(4) (2010) 576–89.

[63] K. Rummens, S.J. van Cromphaut, G. Carmeliet, E. van Herck, R. van Bree, I. Stockmans, R. Bouillon, J. Verhaeghe, Pregnancy in mice lacking the vitamin D receptor: normal maternal skeletal response, but fetal hypomineralization rescued by maternal calcium supplementation, Pediatr Res 54(4) (2003) 466–73.

[64] S.K. Das, Regional development of uterine decidualization: molecular signaling by Hoxa-10, Mol Reprod Dev 77(5) (2010) 387–96.

[65] H.L. Franco, K.Y. Lee, C.A. Rubel, C.J. Creighton, L.D. White, R.R. Broaddus, M.T. Lewis, J.P. Lydon, J.W. Jeong, F.J. DeMayo, Constitutive activation of smoothened leads to female infertility and altered uterine differentiation in the mouse, Biol Reprod 82(5) (2010) 991–9.

[66] M.B. Meyer, S.M. Lee, A.H. Carlson, N.A. Benkusky, M. Kaufmann, G. Jones, J.W. Pike, A chromatin-based mechanism controls differential regulation of the cytochrome P450 gene Cyp24a1 in renal and non-renal tissues, J Biol Chem 294(39) (2019) 14467–14481.

[67] M.B. Meyer, N.A. Benkusky, M. Kaufmann, S.M. Lee, R.R. Redfield, G. Jones, J.W. Pike, Targeted genomic deletions identify diverse enhancer functions and generate a kidney-specific, endocrine-deficient Cyp27b1 pseudo-null mouse, J Biol Chem 294(24) (2019) 9518–9535.

[68] Q. Li, A. Kannan, A. Das, F.J. Demayo, P.J. Hornsby, S.L. Young, R.N. Taylor, M.K. Bagchi, I.C. Bagchi, WNT4 acts downstream of BMP2 and functions via beta-catenin signaling pathway to regulate human endometrial stromal cell differentiation, Endocrinology 154(1) (2013) 446–57.

[69] H.L. Franco, D. Dai, K.Y. Lee, C.A. Rubel, D. Roop, D. Boerboom, J.W. Jeong, J.P. Lydon, I.C. Bagchi, M.K. Bagchi, F.J. DeMayo, WNT4 is a key regulator of normal postnatal uterine development and progesterone signaling during embryo implantation and decidualization in the mouse, FASEB J 25(4) (2011) 1176–87.

[70] Q. Li, A. Kannan, W. Wang, F.J. Demayo, R.N. Taylor, M.K. Bagchi, I.C. Bagchi, Bone morphogenetic protein 2 functions via a conserved signaling pathway involving Wnt4 to regulate uterine decidualization in the mouse and the human, J Biol Chem 282(43) (2007) 31725–32.

[71] B.A. Benayoun, E.A. Pollina, D. Ucar, S. Mahmoudi, K. Karra, E.D. Wong, K. Devarajan, A.C. Daugherty, A.B. Kundaje, E. Mancini, B.C. Hitz, R. Gupta, T.A. Rando, J.C. Baker, M.P. Snyder, J.M. Cherry, A. Brunet, H3K4me3 breadth is linked to cell identity and transcriptional consistency, Cell 158(3) (2014) 673–88.

[72] M.P. Creyghton, A.W. Cheng, G.G. Welstead, T. Kooistra, B.W. Carey, E.J. Steine, J. Hanna, M.A. Lodato, G.M. Frampton, P.A. Sharp, L.A. Boyer, R.A. Young, R. Jaenisch, Histone H3K27ac separates active from poised enhancers and predicts developmental state, Proc Natl Acad Sci U S A 107(50) (2010) 21931–6.

[73] D. Pham, C.E. Moseley, M. Gao, D. Savic, C.J. Winstead, M. Sun, B.L. Kee, R.M. Myers, C.T. Weaver, R.D. Hatton, Batf Pioneers the Reorganization of Chromatin in Developing Effector T Cells via Ets1-Dependent Recruitment of Ctcf, Cell Rep 29(5) (2019) 1203–1220 e7.

[74] X. Wang, Y. Hong, J. Zou, B. Zhu, C. Jiang, L. Lu, J. Tian, J. Yang, K. Rui, The role of BATF in immune cell differentiation and autoimmune diseases, Biomark Res 13(1) (2025) 22.

[75] M.B. Stope, A. Mustea, N. Sanger, R. Einenkel, Immune Cell Functionality during Decidualization and Potential Clinical Application, Life (Basel) 13(5) (2023).

[76] I. Granot, Y. Gnainsky, N. Dekel, Endometrial inflammation and effect on implantation improvement and pregnancy outcome, Reproduction 144(6) (2012) 661–8.

[77] I. Orlov, N. Rochel, D. Moras, B.P. Klaholz, Structure of the full human RXR/VDR nuclear receptor heterodimer complex with its DR3 target DNA, EMBO J 31(2) (2012) 291–300.

[78] P.L. Shaffer, D.T. Gewirth, Structural basis of VDR-DNA interactions on direct repeat response elements, EMBO J 21(9) (2002) 2242–52.

[79] J. Zheng, M.R. Chang, R.E. Stites, Y. Wang, J.B. Bruning, B.D. Pascal, S.J. Novick, R.D. Garcia-Ordonez, K.R. Stayrook, M.J. Chalmers, J.A. Dodge, P.R. Griffin, HDX reveals the conformational dynamics of DNA sequence specific VDR co-activator interactions, Nat Commun 8(1) (2017) 923.

[80] H. Ashour, S.M. Gamal, N.B. Sadek, L.A. Rashed, R.E. Hussein, S.S. Kamar, H. Ateyya, M.N. Mehesen, A.M. ShamsEldeen, Vitamin D Supplementation Improves Uterine Receptivity in a Rat Model of Vitamin D Deficiency: A Possible Role of HOXA-10/FKBP52 Axis, Front Physiol 12 (2021) 744548.

[81] M.C. Chien, C.Y. Huang, J.H. Wang, C.L. Shih, P. Wu, Effects of vitamin D in pregnancy on maternal and offspring health-related outcomes: An umbrella review of systematic review and meta-analyses, Nutr Diabetes 14(1) (2024) 35.

[82] H. Liu, Y. Wen, X. Liang, Y. Xu, D. Qiao, C. Yang, M. Han, H. Li, T. Ren, X. Zhang, G. Li, Z. Liu, Prefrontal cortex neural activity predicts reduction of non-suicidal self-injury in adolescents with major depressive disorder: An event related potential study, Front Neurosci 16 (2022) 972870.

[83] R.M. Marquardt, T.H. Kim, J.H. Shin, J.W. Jeong, Progesterone and Estrogen Signaling in the Endometrium: What Goes Wrong in Endometriosis?, Int J Mol Sci 20(15) (2019).

[84] K. Yu, Z.Y. Huang, X.L. Xu, J. Li, X.W. Fu, S.L. Deng, Estrogen Receptor Function: Impact on the Human Endometrium, Front Endocrinol (Lausanne) 13 (2022) 827724.

[85] R. Usategui-Martin, D.A. De Luis-Roman, J.M. Fernandez-Gomez, M. Ruiz-Mambrilla, J.L. Perez-Castrillon, Vitamin D Receptor (VDR) Gene Polymorphisms Modify the Response to Vitamin D Supplementation: A Systematic Review and Meta-Analysis, Nutrients 14(2) (2022).

