## Supplementary Methods and Figures for "Impact of vitamin D deficiency on defective endometrial decidualization and the repressive role of vitamin D receptor (VDR) in the epigenomic network"

**Author ORCIDs**

MyeongJin Yi https://orcid.org/0000-0002-6561-3620; Skylar G. Montague Redecke https://orcid.org/0000-0002-1418-8863; Tianyuan Wang https://orcid.org/0000-0002-3970-0771; Austin Bell-Hensley https://orcid.org/0000-0003-1498-3012; Shuyun Li https://orcid.org/0000-0002-6172-784X; Abdull J. Massri https://orcid.org/0000-0003-1314-6650; Anne Marie Z. Jukic https://orcid.org/0000-0001-8635-9505; Francesco J. DeMayo https://orcid.org/0000-0002-9480-7336

*Corresponding authors

Correspondence to: National Institute of Environmental Health Sciences, National Institutes of Health, Research Triangle Park, North Carolina, 27709, United States.

**List of Supplementary Data**

**Supplementary methods**

**Blood collection and serum hormone analysis by enzyme-linked immunosorbent assay (ELISA)**

To confirm the level of total serum 25-hydroxyvitamin D (25(OH)D) reaches a deficient status, each blood sample was collected at the terminal point via cardiac puncture. After blood was collected into a yellow cap serum tube, the samples were centrifuged. The upper component (serum) was transferred to a clear tube and maintained on ice during the procedure. An ELISA was adapted to quantify the total serum 25(OH)D level by measuring the bound monoclonal antibody to 25(OH)D. The serum 25(OH)D level was examined using an enzyme immunoassay with a Mouse/Rat 25-OH Vitamin D ELISA Assay kit, according to the manufacturer's instructions (Eagle Biosciences).

**Observation of vaginal smear cytology**

To monitor estrous cycle stages, daily vaginal cytology was performed on female mice that had been maintained on vitamin D-sufficient or -deficient diets for 8 weeks. Mice were 6 weeks old at the start of dietary intervention to avoid puberty-related confounding factors. Vaginal cytology screening was performed after 8 weeks on the diet and continued for 15 consecutive days (corresponding to weeks 8–10 of the diet). Briefly, 10 μL of sterile PBS was gently pipetted into the vaginal opening and aspirated to collect cells. The specimen was transferred to a glass slide, air-dried, and examined under a light microscope. Vaginal cell types were classified as leukocytes, cornified epithelial cells, or nucleated epithelial cells [1]. The stage of the estrous cycle, proestrus, estrus, metestrus, or diestrus, was determined based on the relative proportion of cell types [1]. Healthy female mice typically exhibit 4–5-day estrous cycles, and daily monitoring over 15 days allowed observation of multiple full cycles per mouse [1].

**Superovulation**

Similar to the vaginal smear cytology, mice were 6 weeks old at the start of the dietary intervention to avoid puberty-related confounding factors. At the collection stage, female mice (14–15 weeks old) that had been maintained on a vitamin D-sufficient (n = 3) or -deficient diet (n = 4) for 8 weeks were subjected to a superovulation protocol. Mice were intraperitoneally injected with 3.25 IU equine chorionic gonadotropin (eCG), followed by 2 IU human chorionic gonadotropin (hCG) 45 h later [2]. At 21 h post-hCG injection, oviducts were collected and flushed for oocyte counting.

**Breeding trial**

A six-month breeding trial was performed to assess the fertility of female mice maintained on vitamin D-sufficient or -deficient diets. Female mice (6 weeks old) were housed on their assigned diets throughout the trial and paired with fertile males (8 weeks old) daily. To minimize the dietary influence on males, pairing was performed each evening at 17:00, and males were separated the following morning at 09:00. Vaginal plugs were checked each morning to monitor mating. This pairing/separation cycle was repeated for 6 months, and females remained on their assigned diets for the entire duration of the study.

At study initiation, seven females were assigned to each diet group (n = 7 per group) and were paired nightly with fertile males as described above. During the trial, three females in the vitamin D-deficient cohort died at delivery of their first litters. To maintain breeding throughput, three age-matched replacement females (same strain and supplier) were introduced into the deficient cohort and maintained on the assigned diet for the remainder of the study (resulting in a total of 10 females enrolled in the deficient arm over the course of the trial). Because these replacement animals entered the study after it began and therefore had a shorter cumulative exposure to the assigned diet, all primary fertility indices and comparative analyses were calculated using the original seven females per group (i.e., the animals present at study initiation). Replacement animals were retained for colony continuity but were excluded from the primary fertility endpoint calculations to avoid bias introduced by post-randomization replacement and unequal diet exposure.

**Validation of VDR binding motifs and peak reliability in endometrial stromal cells using kidney-derived data**

Even though numerous studies have demonstrated that endometrial stromal cells express VDR, its relatively low expression impeded our cistromic analysis. To overcome this issue, we activated VDR using a lentiviral vector. However, concerns regarding the reliability and validation of this data prompted us to compare our cistromic results to publicly available data sets derived from the kidney cortex of a deceased 69-year-old female donor, obtained through the University of Wisconsin-Madison Organ Procurement Program [3, 4]. We utilized technical triplicates (GFM3716705, GFM3716706, and GFM3716707) from dataset GSE129585 [3, 4]. Since the original coordinates were based on the hg19 genome assembly, we converted coordinates to hg38 using the UCSC liftOver tool. To eliminate potential false positives, peaks on chromosome Y were excluded, as the kidney cortex originated from a female donor.

**Supplementary Figure 1. Assessment of vitamin D deficiency diet studies in C57BL/6J female mice.** (A) Screening of total serum 25(OH)D levels in mice after dietary intervention from 5 weeks to 18 weeks of each diet. (B) Schematic illustration of the vitamin D deficiency diet protocol in the estrous cycle and superovulation, and (C) in a 6-month breeding trial.

**Supplementary Figure 2. Fertility outcomes in vitamin D-deficient mice.** (A) Analysis of estrous cycle progression over 15 days in vitamin D-control and vitamin D-deficient mice. (B) Time point of superovulation and number of oocytes collected following superovulation induction. (C) Fertility indices, including the total number of litters and pups in each group. (D) Cumulative pup counts over the breeding trial duration. (E) Incidence of dystocia-related mortality in vitamin D-deficient dams. Data are presented as mean ± SEM.

**Supplementary Figure 3. Summary of RNA bulk sequencing in T-HESCs by VDR knockdown and 1,25(OH)$_2$D$_3$.** (A) Principal component analysis (PCA) illustrating distinct transcriptomic profiles in T-HESCs treated with control siRNA (siNT) or VDR-targeting siRNA (siVDR), and either vehicle or 2 nM 1,25(OH)$_2$D$_3$ for 24 hours. PCA highlights the separation of samples based on transcriptomic shifts induced by VDR knockdown and 1,25$(OH)_2D_3$ treatment. (B) Heatmap of differentially expressed genes (DEGs) across experimental conditions (siNT or siVDR, with vehicle or 1,25$(OH)_2D_3$). Hierarchical clustering demonstrates the transcriptomic impact of VDR knockdown and 1,25$(OH)_2D_3$ on gene expression profiles. (C) Summary of DEG analysis. Bar graph showing the number of upregulated and downregulated DEGs in siNT vs. siVDR cells under vehicle or 1,25$(OH)_2D_3$ treatments. The data illustrate the substantial transcriptomic influence of VDR and its ligand. (D) Chromatin accessibility analysis: 19,189 genes were annotated in open chromatin regions (ATAC-seq). Of these, 1,208 out of 1,499 receptor DEGs (80.59%) and 1,802 out of 2,311 ligand-receptor DEGs (77.97%) overlapped with accessible regions. (E) Validation of RNA-seq findings by qPCR analysis of selected genes, including CYP24A1, IGFBP1, EFL1, and VEGFA. CYP24A1 induction by 1,25$(OH)_2D_3$ is attenuated in siVDR cells. VDR knockdown enhances IGFBP1 expression, while the expression of EFL1 and VEGFA is shown to be modulated by 1,25$(OH)_2D_3$ in a VDR-dependent manner.

**Supplementary Figure 4. Pathways regulated by transcriptomic changes in T-HESCs by ligand-dependent DEGs.** Canonical pathways in transcriptomic changes by the 900 overlapped regulated transcriptomes between the receptor (siNT vehicle vs. siVDR vehicle) and the ligand (siNT 1,25$(OH)_2D_3$ vs. siVDR 1,25$(OH)_2D_3$).

**Supplementary Figure 5. Validation of endometrial stromal cell VDR peaks using kidney-derived ChIP-seq data.** (A) HOMER motif analysis of kidney-derived peaks (samples GFM3716705, GFM3716706, and GFM3716707) revealed strong enrichment for VDR binding motifs and renal regulators such as HNF1B. (B) Comparison of peak distributions of human endometrial stromal cells and kidney. Genomic annotation of 30,113 kidney-derived peaks showed a predominant distribution in promoter regions (33.59%). (C) Overlap Venn diagram between kidney peaks and vehicle-treated (314 peaks; 35.48%) or 1,25$(OH)_2D_3$-treated (1,966 peaks; 32.27%) endometrial stromal cell peaks confirmed shared binding regions. Heatmap and metaplot analysis demonstrated increased peak intensity in 1,25$(OH)_2D_3$-treated cells for kidney-overlapping peaks.

**Supplementary Figure 6. Transcriptomic enrichment of chromatin-related receptor genes regulated by VDR in T-HESCs.** (A) List of 27 differentially expressed chromatin-related receptor genes identified from the total set of 1,499 VDR-dependent DEGs. (B) Gene set enrichment analysis (GSEA) bubble plot of enriched Gene Ontology (GO) Biological Process terms associated with chromatin regulation. Bubble size indicates the gene ratio, and color represents the adjusted *P* value. (C) Representative GSEA enrichment plot showing chromatin-related GO term enrichment, with "Rank in ordered dataset" on the *x*-axis and the running enrichment score on the *y*-axis. (D) Summary of chromatin-related GO terms grouped into two major pathways based on normalized enrichment score (NES). Bar color corresponds to the adjusted *P* value, and NES is plotted on the *x*-axis.

**Supplementary Table 1.** Key sources, information on the nutrition of vitamin D diets, and a list of abbreviations.

**Supplementary Table 2.** Information on peaks within open chromatin regions in T-HESCs, both siNT and siVDR.

**Supplementary Table 3.** List of DEGs, canonical pathways, and upstream regulators in T-HESCs regulated by ligand (1,25$(OH)_2D_3$) and receptor (VDR).

**Supplementary Table 4.** Information on peaks and motifs in lentiviral transduced T-HESCs treated with either vehicle or 1,25$(OH)_2D_3$.

**Supplementary Table 5.** Information on peaks and motifs in the human kidney (GSE129585) and overlay peaks and motifs between human kidney and lentiviral transduced T-HESCs treated with either vehicle or 1,25$(OH)_2D_3$.

**Supplementary Table 6.** List of overlapped peaks, DEGs, and motifs between cistromic and transcriptomic analyses.

**Supplementary Figure 1.**

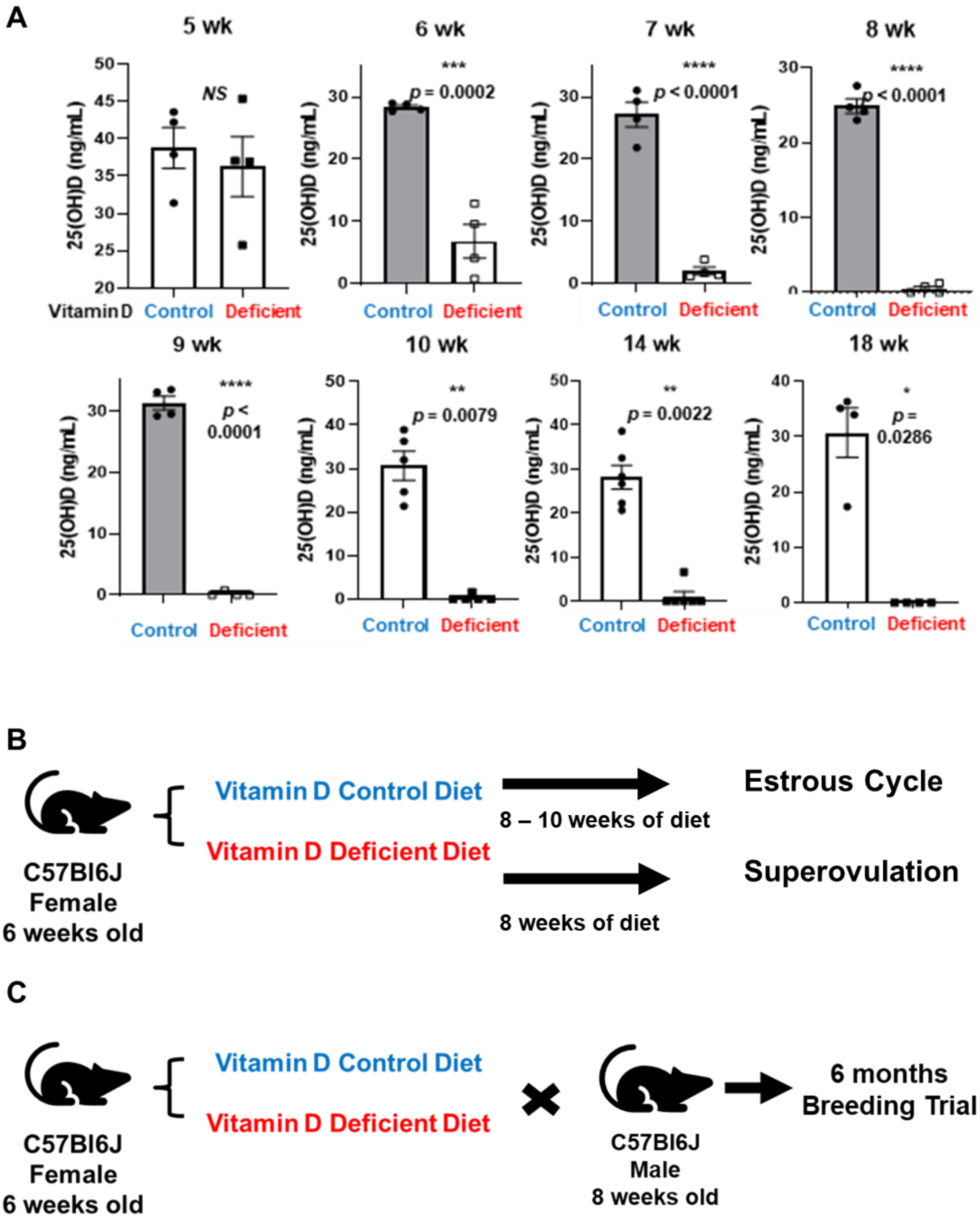

**Supplementary Figure 2.**

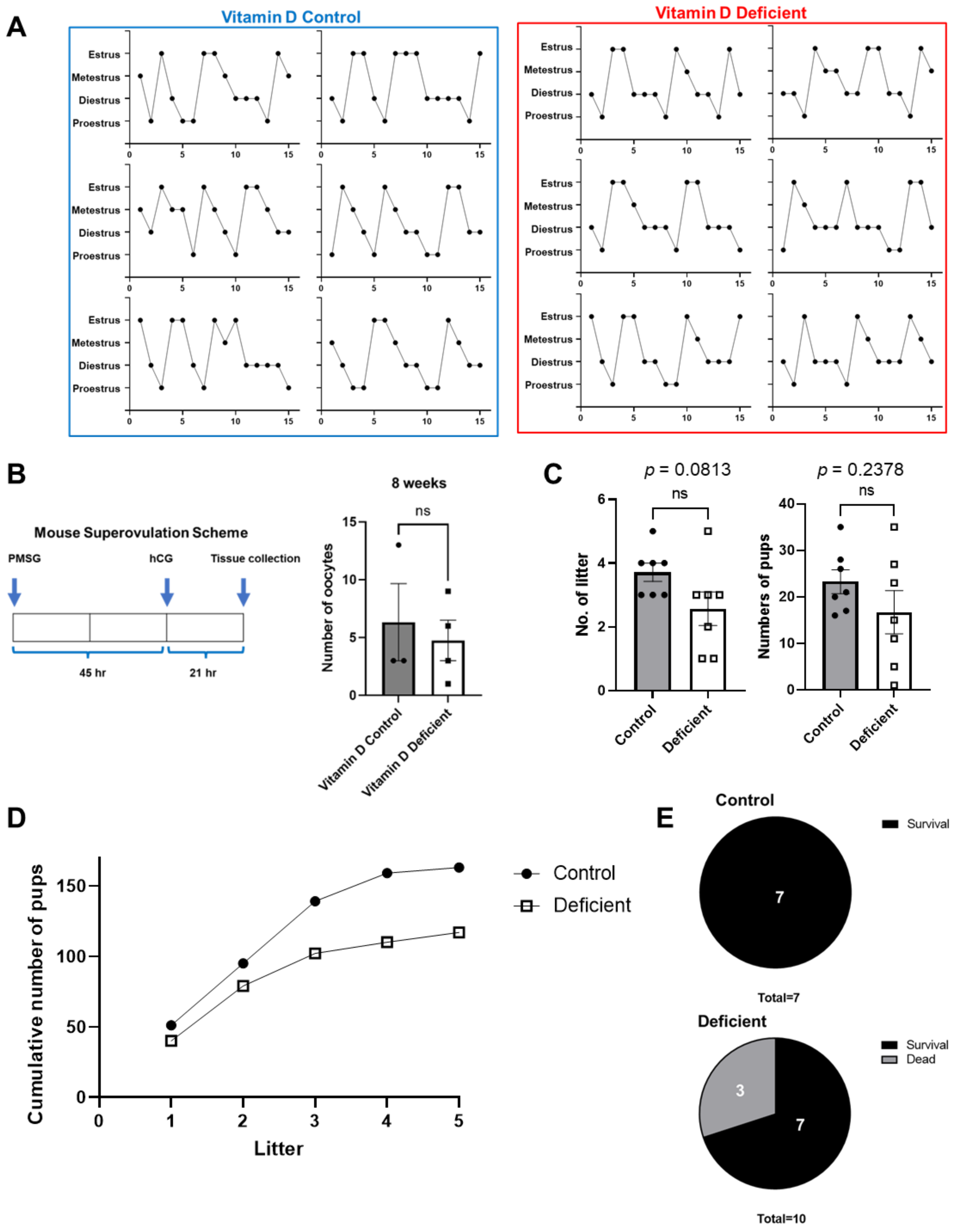

**Supplementary Figure 3.**

A

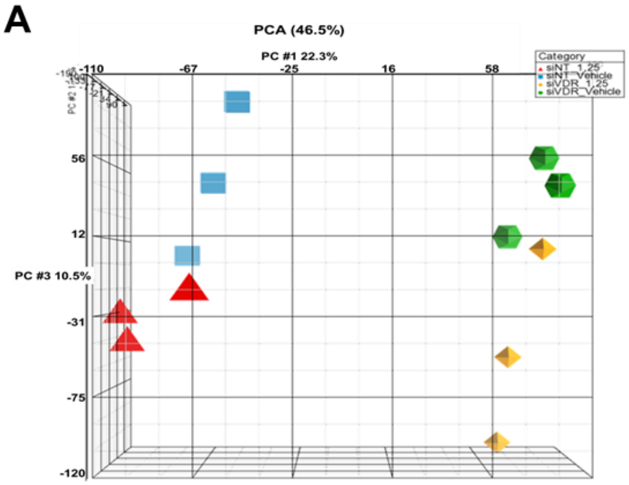


B

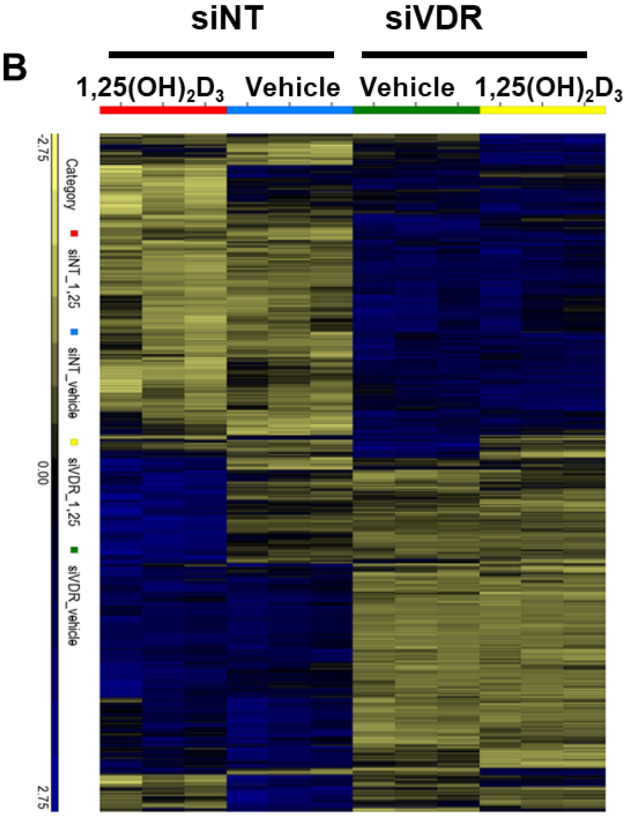


C

| Contrast | FC ≥ \|1.4\| |
|---|---|
| siNT vehicle vs. siVDR vehicle | 1499 (Up: 818, Down: 681) |
| siNT 1,25(OH)$_2$D$_3$ vs. siVDR 1,25(OH)$_2$D$_3$ | 2311 (Up: 1362, Down: 949) |
| siNT 1,25(OH)$_2$D$_3$ vs. siNT vehicle | 626 (Up: 386, Down: 240) |
| siVDR 1,25(OH)$_2$D$_3$ vs. siVDR vehicle | 56 (Up: 8, Down: 48) |

D

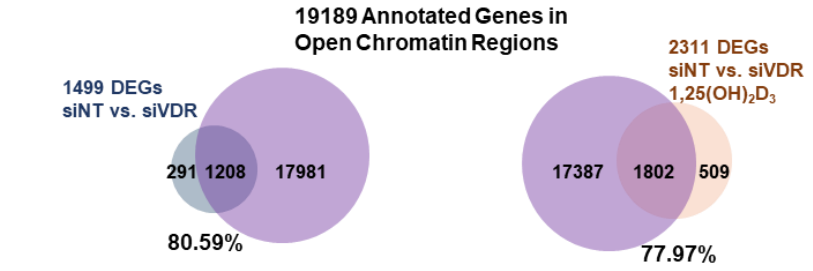


E

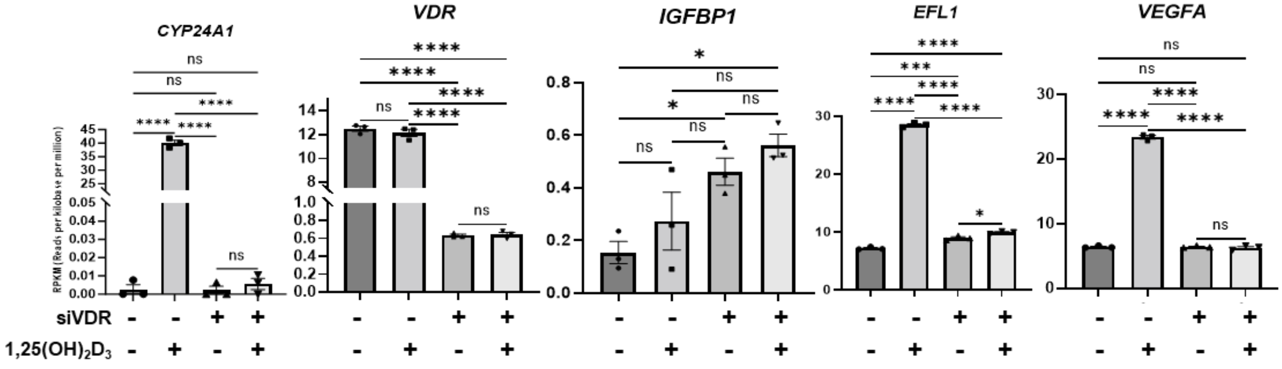

**Supplementary Figure 4.**

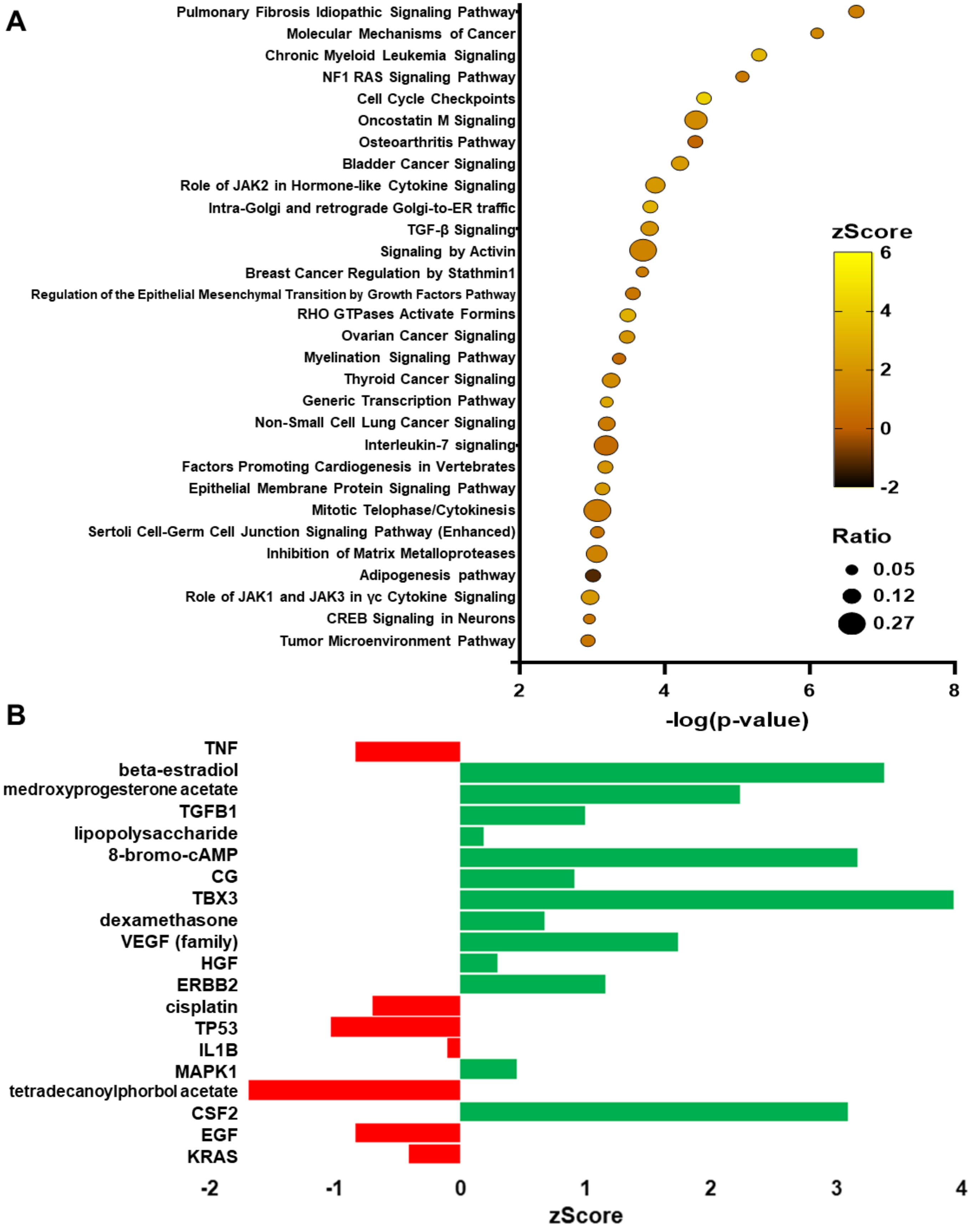

**Supplementary Figure 5.**

**A**

*De novo* motif

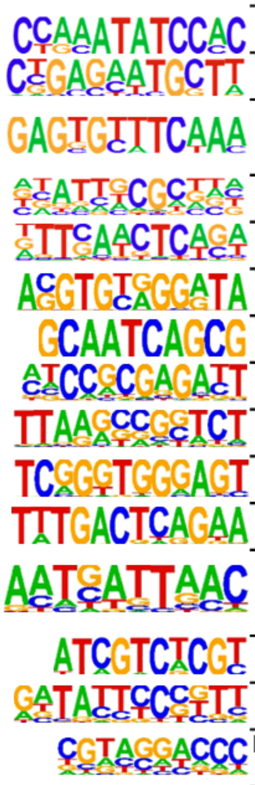

| De novo motif | Best Match | P-value | % of targets |
|---|---|---|---|
| | **ZNF354C**/MA0130.1 | 1e-601 | 0.98% |
| | PB0138.1_**Irf4**_2 | 1e-582 | 1.09% |
| | **CHR**(?)/Hela-CellCycle-Expression | 1e-553 | 1.05% |
| | **CEBPG**/MA0838.1 | 1e-551 | 1.04% |
| | **VDR**/MA0693.2 | 1e-510 | 3.27% |
| | **ZNF75D**/MA1601.1 | 1e-489 | 1.16% |
| | **Dux**/MA0611.1 | 1e-477 | 1.59% |
| | **ZBTB33**/MA0527.1 | 1e-473 | 1.60% |
| | PB0048.1_**Nkx3-1**_1 | 1e-473 | 1.07% |
| | PB0167.1_**Sox13**_2 | 1e-473 | 1.55% |
| | **BATF3**/MA0835.2 | 1e-432 | 1.29% |
| | **HNF1b**(Homeobox)/PDAC-HNF1B-ChIP-Seq | 1e-425 | 6.92% |
| | **TFEC**/MA0871.2 | 1e-422 | 1.03% |
| | PH0126.1_**Obox6** | 1e-418 | 1.19% |
| | **LRF**(Zf)/Erythroblasts-ZBTB7A-ChIP-Seq | 1e-407 | 1.53% |

**B**

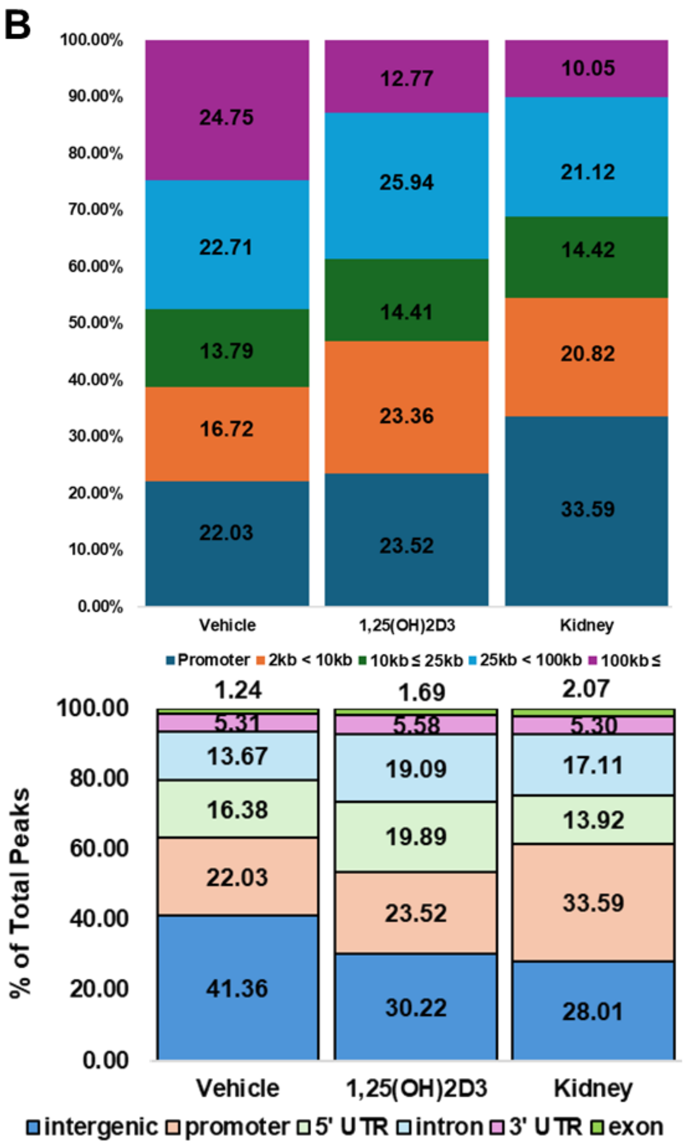


**C**

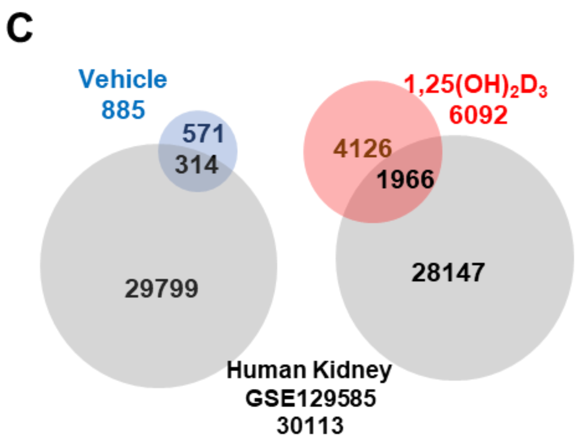


**35.48% of vehicle-treated and 32.27% of 1,25(OH)2D3-treated human endometrial stromal cell peaks overlapped with human kidney peaks**

**Supplementary Figure 6.**

**A**

| Chromatin-related Receptor DEGs |
|---|
| PADI2, MYOCD, MEN1, N6AMT1, LSM11, DCP2, HJURP, SNAI1, MECOM, H2AJ, LMNB2, SETD7, LMNB1, RB1, SIN3A, BCL6, MYBL1, EID3, MKI67, HMGA1, H2BC4, SOX9, KAT6B, RSF1, NAP1L3, ESCO2, PLK1 |

**B**

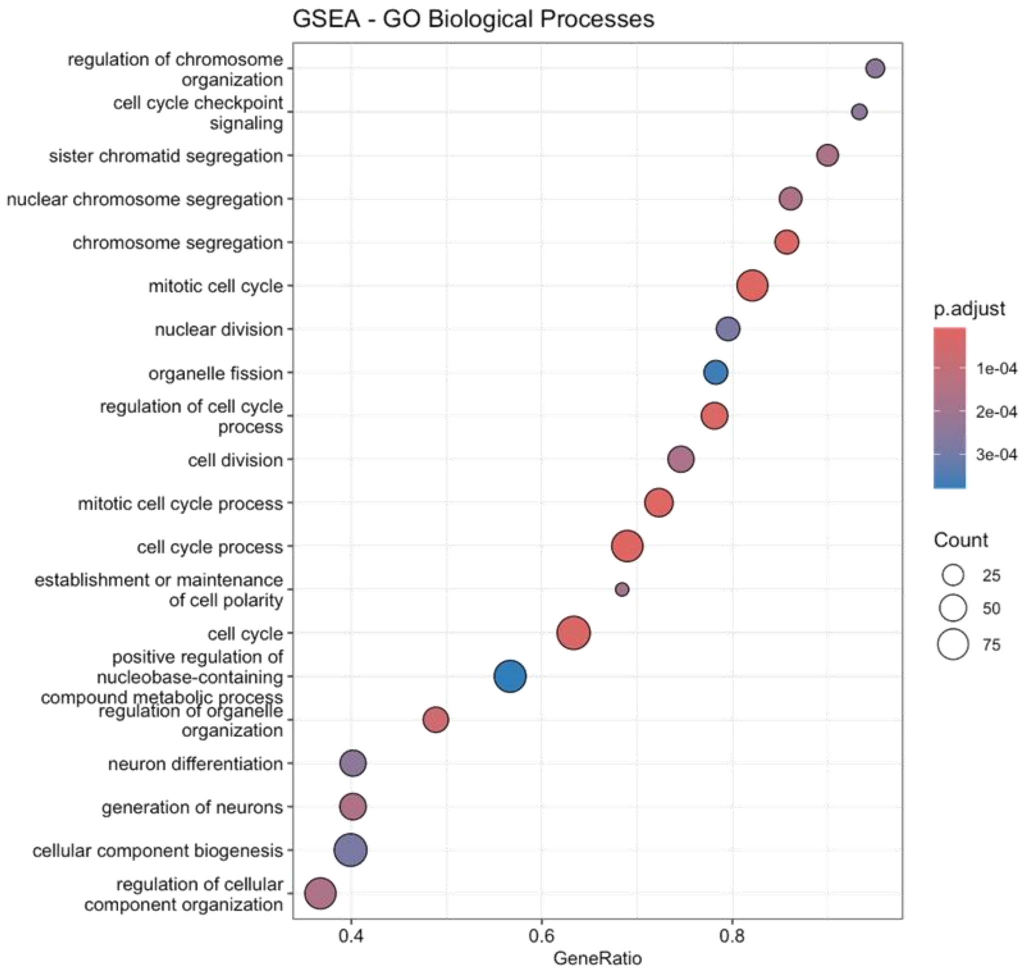


**C** **D**

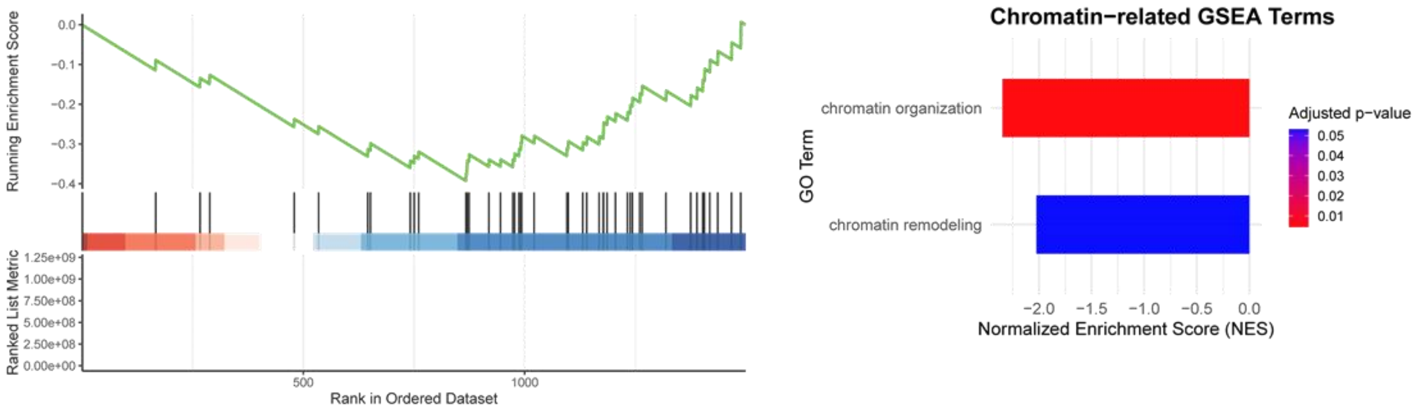

### References

[1] S.L. Byers, M.V. Wiles, S.L. Dunn, R.A. Taft, Mouse estrous cycle identification tool and images, PLoS One 7(4) (2012) e35538.

[2] R. Li, S.P. Wu, L. Zhou, B. Nicol, J.P. Lydon, H.H. Yao, F.J. DeMayo, Increased FOXL2 expression alters uterine structures and functions, Biol Reprod 103(5) (2020) 951-965.

[3] M.B. Meyer, S.M. Lee, A.H. Carlson, N.A. Benkusky, M. Kaufmann, G. Jones, J.W. Pike, A chromatin-based mechanism controls differential regulation of the cytochrome P450 gene Cyp24a1 in renal and non-renal tissues, J Biol Chem 294(39) (2019) 14467-14481.

[4] M.B. Meyer, N.A. Benkusky, M. Kaufmann, S.M. Lee, R.R. Redfield, G. Jones, J.W. Pike, Targeted genomic deletions identify diverse enhancer functions and generate a kidney-specific, endocrine-deficient Cyp27b1 pseudo-null mouse, J Biol Chem 294(24) (2019) 9518-9535.
